# Local sympathetic neurons regulate adaptive immunity in response to Streptococcus pneumoniae infection by modulating T- and B-cell effector functions

**DOI:** 10.64898/2026.02.16.706205

**Authors:** Fengli Zhu, Kayla Davis, Sreemoyee Acharya, Rishit Pattanashetti, Jinendiran Sekar, Diane Aguilar, Nathisha Kalpage, Omid Akbari, Srividya Swaminathan, Peter Jorth, Marc Swidergall, Nicholas Jendzjowsky

## Abstract

Adaptive immunity in the lung must be rapidly mobilized to control bacterial pathogens, yet the neuroimmune signals that shape local B and T cell responses remain poorly defined. Here we identify a sympathetic–immune axis that regulates lung-resident memory B cells, antigen-specific IgG production, and protection against *Streptococcus pneumoniae*. Lung-targeted chemical sympathectomy reduced norepinephrine availability, diminished Tbet⁺ memory and resident-memory B cell populations, and markedly impaired *S. pneumoniae*–specific IgG, resulting in increased bacterial burden. Mice lacking β₁- and β₂-adrenergic receptors phenocopied these defects, and adoptive transfer of *Adrb1^⁻/⁻^Adrb2^⁻/⁻^* B cells into *Ighm^⁻/⁻^* hosts was sufficient to reduce IgG and worsen infection. Combined T cell depletion and B cell–intrinsic adrenergic receptor loss further impaired bacterial control and suppressed IFNγ, indicating that norepinephrine stimulates T cell IFNγ release and that norepinephrine and IFNγ together drive optimal B cell IgG production. In vitro, norepinephrine enhanced T cell IFNγ secretion, and norepinephrine plus IFNγ synergistically increased IgG. These findings define norepinephrine as a key regulator of lung-resident humoral immunity and reveal a neuroimmune mechanism that promotes antibody-mediated bacterial clearance at the respiratory mucosa.

**Graphical abstract:** 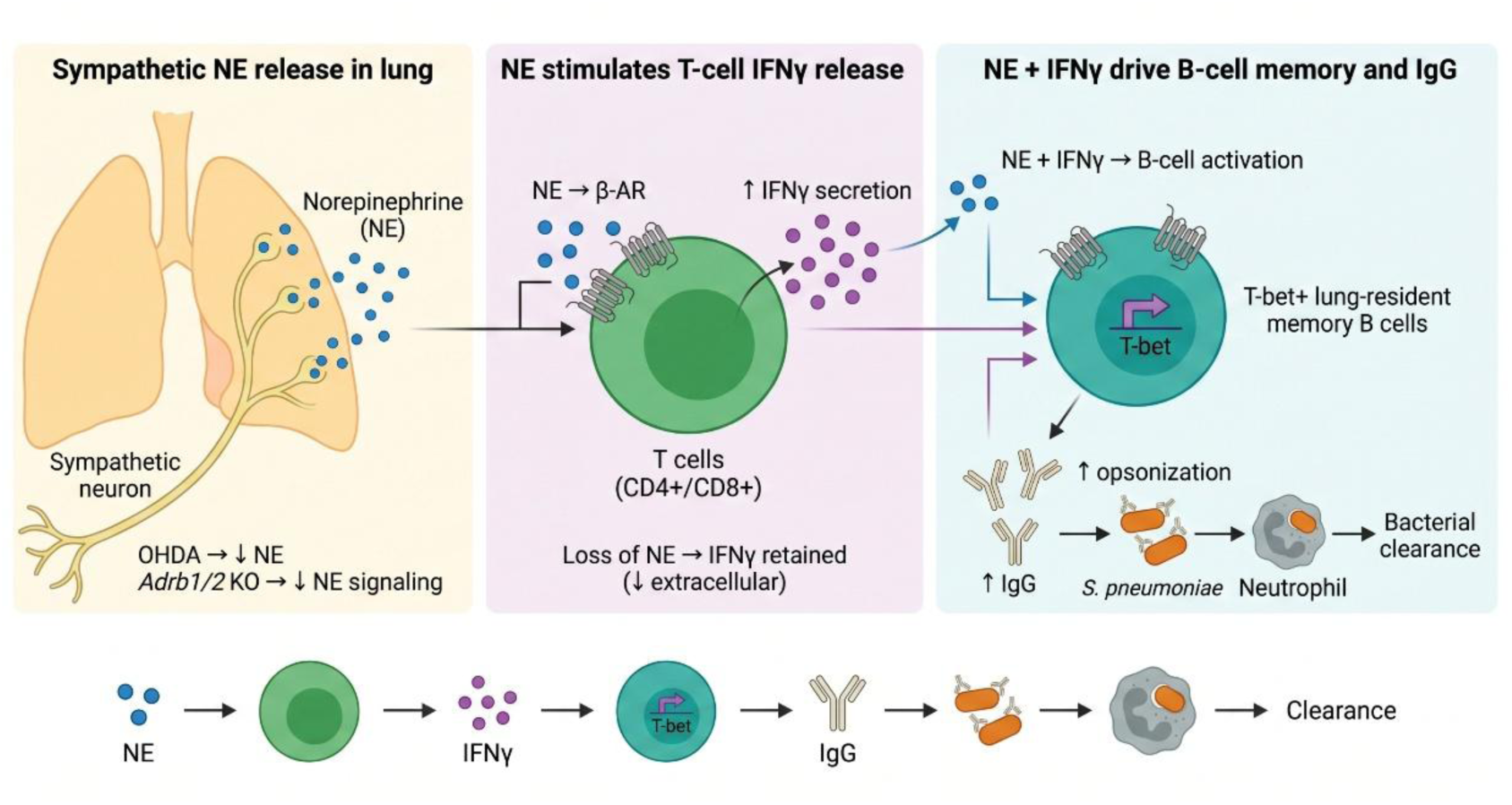

**Highlights:** Our results identify norepinephrine as a key regulator of lung-resident memory B cell differentiation and IgG-mediated bacterial clearance. By linking sympathetic signaling to T cell IFNγ release and B cell activation, this work reveals a neuroimmune axis that shapes mucosal humoral immunity. Targeting this pathway may enhance protection against respiratory bacterial infections.

## Introduction

The neuroimmune interplay in the lung profoundly affects innate and adaptive immunity during bacterial infection(**1**). Stimulation of sensory neurons by cytokines(**2–6**), proteases(**7–11**), pathogen-associated molecular patterns(**12–14**) and immunoglobulins(**15–18**) lead to neuronal depolarization and the release of neuropeptides into the immediate vicinity to assist with immune coordination(**19, 20**). Such sensory neuropeptide release has been demonstrated to stimulate T cells(**13**), and B cells(**21**), having a determinate effect on cell effector function in a disease-specific context(**1**).

Classic reflex arcs, which integrate signals in the brainstem and subsequently stimulate efferent nerves to release neurotransmitters within the target tissue, are activated by sensory neurons and provide an additional neuroimmune mechanism for immune modulation. Within the lung, a majority of classic sensory reflexes stimulate efferent pathways within the vagus(**22**), which is considered to be primarily of parasympathetic origin. Recent studies have delineated a population of tyrosine hydroxylase-positive neurons that are likely to be of sympathetic lineage(**23–25**). Chemical sympathectomy in the lungs halted innate immunity in response to acute LPS treatment and exacerbated inflammation(**24**) whereas vagal sensory stimulation during acute viral stimulation reduced inflammation via a systemic sympathetic circuit involving the adrenal medulla(**26**).

We have recently demonstrated that the release of specific neuropeptides by sensory neurons profoundly affects humoral immunity(**21**) via the stimulation of immunoglobulin production by B lineage cells. Given that a near-simultaneous stimulation of sensory and efferent neurons occurs in response to sensory nerve stimulation, we asked whether sympathetic efferent neurons would contribute to adaptive immunity in response to *Streptococcus pneumoniae* pre-exposure and infection, given that norepinephrine has been shown to affect IgG production(**27**).

We determined that efferent tyrosine hydroxylase^+^ neurons are activated in response to *S. pneumoniae*. We showed that suppression of norepinephrine release via chemical sympathectomy hinders the development of B memory and B resident memory cells, *S. pneumoniae antigen*-specific IgG release, and worsens bacterial burden. Sympathetic depletion prevented T cell IFNγ release, resulting in a reduced concentration in the lungs. Deletion of β-adrenergic receptors phenocopied these effects, and this held true only for B cell receptor deletion. When we depleted CD4^+^ and CD8^+^ T cells and deleted the β-adrenergic receptor on B cells, we observed decreased IFNγ and worsened disease burden. *In vitro* experiments demonstrated that norepinephrine stimulates T cell IFNγ release and that norepinephrine and IFNγ stimulate B cell IgG release. Finally, intact adrenergic receptors on B cells are required to produce adequate opsonization and bacterial killing

## Results

### Lung-targeted sympathetic knockdown exacerbates lung infection

Global sympathetic suppression may ablate either a systemic circuit involving various postganglionic sympathetic nerves or the adrenal medulla(**26**) in addition to lung-innervating sympathetic neurons(**23, 24**). Therefore, to target the lung-innervating tyrosine-hydroxylase+ (Th+) nerve fibers, potentially involved in stimulating adaptive immunity, we delivered 6-hydroxydopamine (6-OHDA) intranasally, as was recently completed(**24**), to reduce lung-innervating sympathetic neurons. In response to our model of *S. pneumoniae 19F* pre-exposure and infection (Figure 1A), norepinephrine release was suppressed in intranasally treated 6-OHDA mice compared with vehicle-treated mice and was reduced to a level comparable to that of naïve mice (Figure 1B). However, acetylcholine, the primary parasympathetic neurotransmitter, remained unaffected by sympathetic neuron ablation (Figure 1C). The reduction of norepinephrine release coincided with reduced immunostaining for Th+ neurons in the lungs in distal and proximal airways when mice were treated with intranasal 6-OHDA delivery (Figure 1D, S1, SV1-6). The effect of lung-targeted 6-OHDA was confined to the lungs, as Th+ staining in the aorta, innervated by sympathetic efferent neurons(**28**), was similar between vehicle and 6-OHDA groups, indicating lung-specific effects and few off-target effects of our 6-OHDA lung delivery, consistent with previous results(**24**).

**Figure 1.**
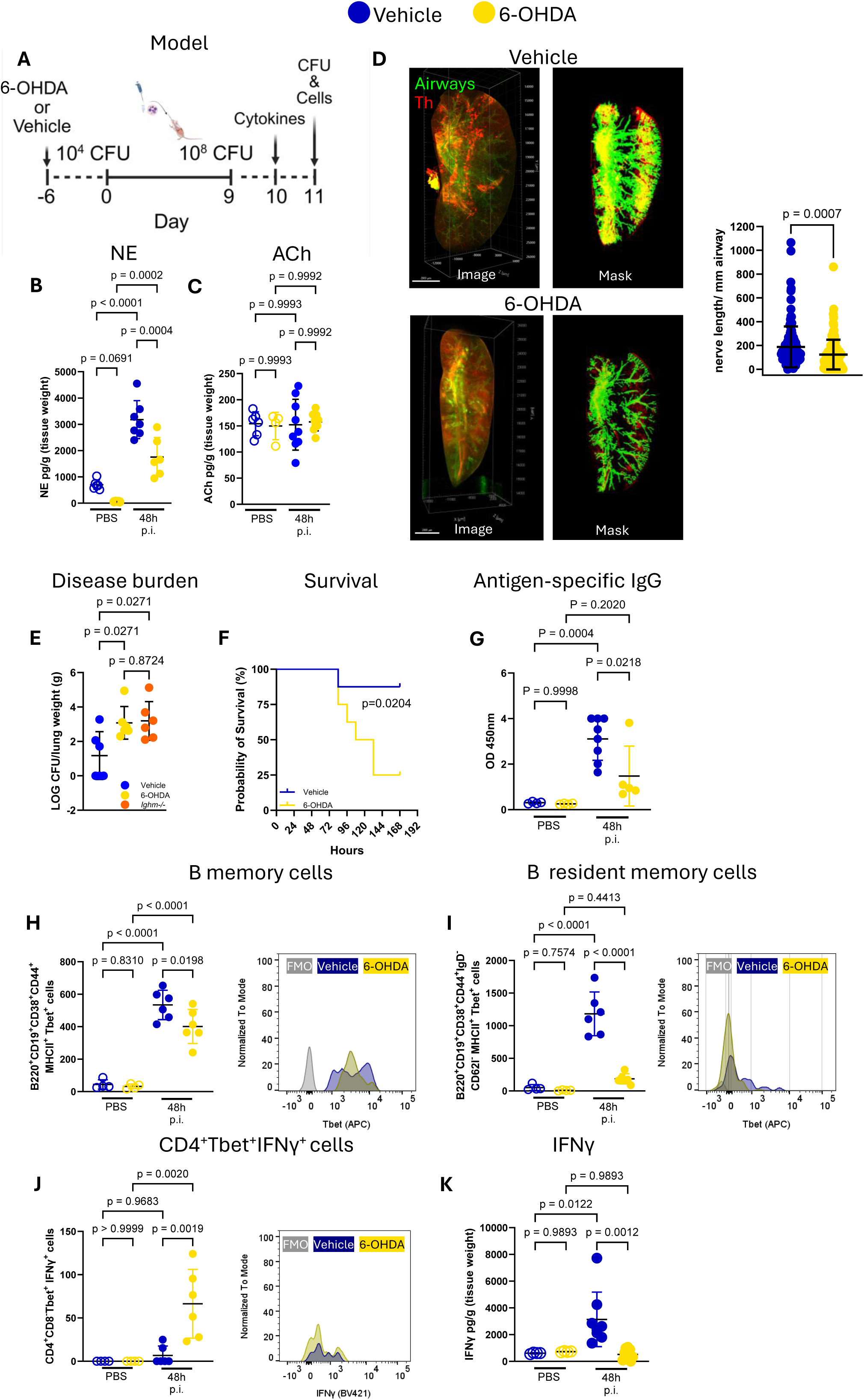
Lung-targeted sympathetic ablation increases disease burden due to a reduction in B cells. A) Mice were randomized to receive vehicle or 6-OHDA and then subjected to *S. pneumoniae* pre-exposure and infection. B) 6-OHDA reduced norepinephrine (NE, primary sympathetic neurotransmitter) release in the lung in response to *S. pneumoniae.* ANOVA with Holm-Sidak post-hoc test; n=4-7. C) 6-OHDA did not alter acetylcholine (ACh, primary parasympathetic neurotransmitter) release in the lung in response to *S pneumoniae.* ANOVA with Holm-Sidak post-hoc test; n=4-9. D) Vehicle and 6-OHDA representative images and computationally derived masks are shown to illustrate the depletion of tyrosine hydroxylase+ (Th) in airways (FITC-streptavidin). Unpaired t-test. N=3 mice, n=40-42 airways per mouse were analyzed. Scale bar = 2000µm. E) Disease burden was greater in 6-OHDA compared to vehicle mice and similar to *Ighm^-/-^* mice. ANOVA with Holm-Sidak post-hoc test; n=6. F) In response to pre-exposure with *S. pneumoniae 19F* and infection with *6303*, 6-OHDA mice had worse survival compared to vehicle mice. Log-rank (Mantel-Cox) test; n=8. G) *S. pneumoniae* antigen-specific IgG was reduced in the lungs of 6-OHDA mice. One-way ANOVA with Holm-Sidak post-hoc test; n=4-8. H) B memory B220^+^ CD19^+^ CD38^+^ CD44^+^ MHCII^+^ Tbet^+^ cells were reduced with 6-OHDA. One-way ANOVA with Holm-Sidak post-hoc test; n=4-8. I) B resident memory B220^+^ CD19^+^ CD38^+^ CD44^+^ IgD^-^ CD62l^-^ MHCII^+^ Tbet^+^ cells were reduced with 6-OHDA. One-way ANOVA with Holm-Sidak post-hoc test; n=4-8. J) CD4^+^ Tbet^+^ IFNγ^+^ cells were increased with 6-OHDA. One-way ANOVA with Holm-Sidak post-hoc test; n=4-8. K) IFNγ concentration in the lungs was reduced with 6-OHDA. One-way ANOVA with Holm-Sidak post-hoc test; n=4-9. Two-three independent experiments for each analysis.

Interestingly, ablation of lung-innervating sympathetic neurons worsened disease burden in response to *S. pneumoniae 19F* pre-exposure and infection (Figure 1E) and survival following *19F* pre-exposure and infection with a lethal *S. pneumoniae* strain *6303* (Figure 1F). To compare disease burden in a model lacking an adaptive immune response, we observed that 6-OHDA-treated mice had disease burden similar to that of *Ighm-/-* mice (which lack mature B cells; Figure 1E). Therefore, we hypothesized that norepinephrine released from sympathetic lung-innervating neurons stimulates an adaptive immune response.

### Sympathetic ablation reduces IgG and lung-resident memory B cells

We first investigated whether *S. pneumoniae* antigen-specific IgG was suppressed with 6-OHDA lung treatment and found that lung-specific sympathetic ablation reduced *S. pneumoniae* antigen-specific IgG titers (Figure 1G). Therefore, we analyzed lung-extravascular B cells, as the lungs were transcardially perfused prior to analysis, and found that total memory-type B cell counts were suppressed by 6-OHDA treatment (Figure 1H, S2, S3). When we further analyzed extravascular B cells, we found that in vehicle-treated, Th^+^ neuron-intact mice, a significantly greater population of memory and resident memory B cells expressed Tbet+, a transcription factor required for the establishment of memory(**29, 30**) were reduced by intranasal 6-OHDA treatment (Figure 1I, S2). Because lungs were perfused prior to digestion, the CD62L^−^ CD38^+^ CD44^+^ Tbet^+^ population likely represents extravascular, tissue-resident B cells rather than circulating memory cells. The reduction in this resident-memory population closely paralleled the reduction in *S. pneumoniae–*specific IgG, consistent with their role as rapid extrafollicular antibody producers. Interestingly, plasma cells and isotype-switched memory B cells were not affected by 6-OHDA (Figure S3). We suggest that norepinephrine enhances memory B cell activation to increase the release of antigen-specific IgG(**31**). Interestingly, neutrophil numbers increased in 6-OHDA mice compared with vehicle controls (Figures S2 and S3), suggesting that increased neutrophil numbers cannot overcome a defect in opsonization.

The number of CD4 and CD8 IFNγ-expressing T cells was elevated in 6-OHDA mice (Figure 1J, S2, S3). Although norepinephrine has previously been shown to raise IFNγ levels in T cells(**32**), analysis of lung homogenates for IFNγ release showed that 6-OHDA treatment reduced the amount of available IFNγ (Figure 1K). The increase in intracellular IFNγ+ T cells despite reduced extracellular IFNγ suggests impaired secretion rather than impaired production, consistent with norepinephrine acting as a stimulating signal for IFNγ release. These data suggest that norepinephrine assists with IFNγ secretion rather than IFNγ production, revealing a potential NE-dependent checkpoint in T cell effector function.

### Deletion of adrenergic receptors phenocopies chemical sympathectomy

Norepinephrine binds to α- and β-adrenergic receptors. Using publicly available RNA-seq data from the Immgen database(**33, 34**), the cell types with the most prominent norepinephrine-binding receptors are shown (Figure 2A). Both β-adrenergic 1 and 2 receptors are expressed on lymphocytes, with relatively few nicotinic receptors or acetylcholine precursors; muscarinic receptors appeared to be sparsely represented(**33, 34**) (precursors and receptors for parasympathetic signaling). However, the overwhelmingly predominant receptor expressed on T and B cells is the β2-adrenergic receptor(**35, 36**).

**Figure 2.**
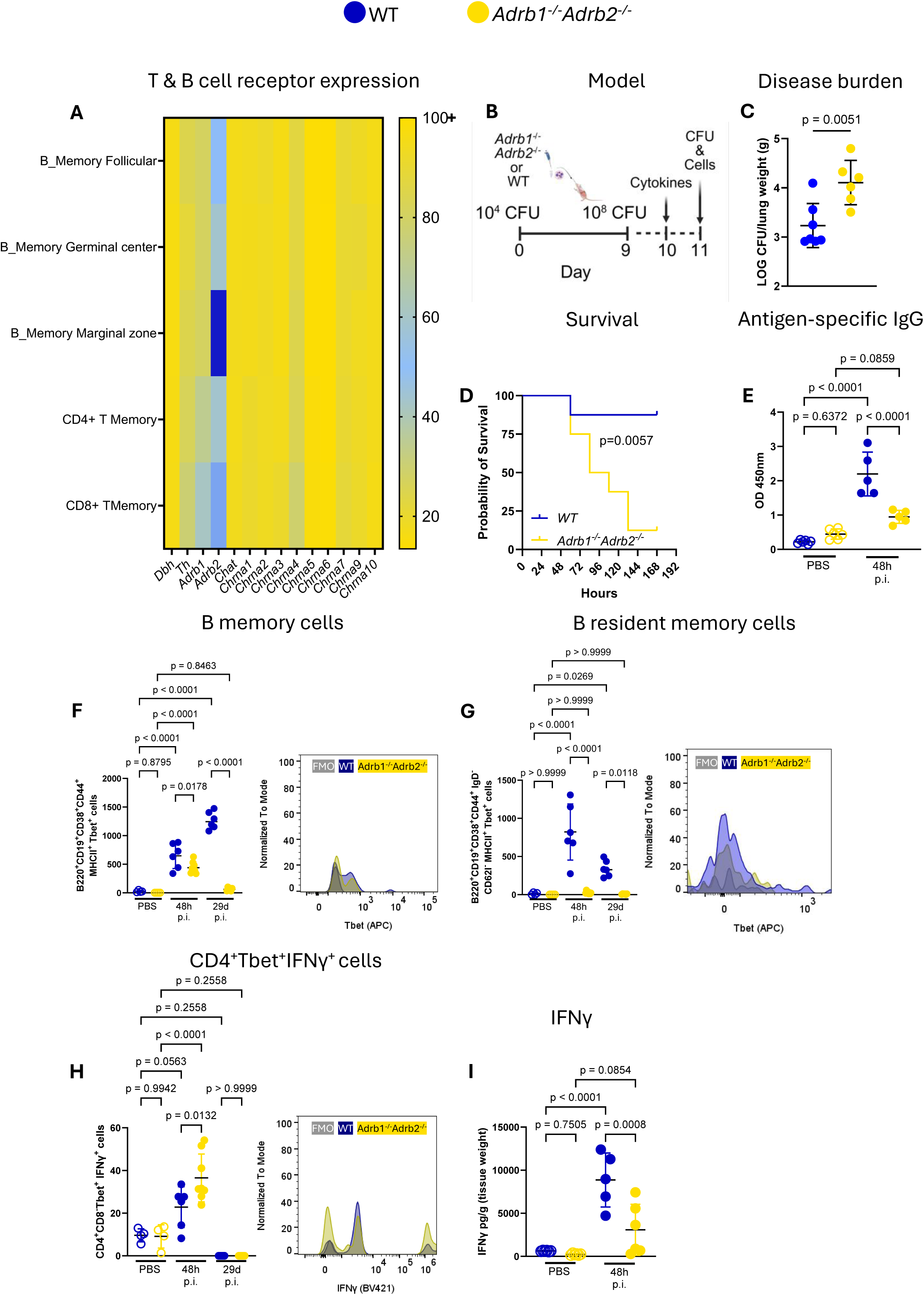
*Adrb1^-/-^Adrb2^-/-^*deletion increases disease burden by reducing B cells 48 h and 29 days post-infection. A) The Immgen database was used to identify prominent adrenergic (*Adrb1, Adrb2*) and nicotinic receptors (*Chrna1-10*) and norepinephrine (*Dbh*- dopamine beta carboxylase, *Th*- tyrosine hydroxylase) and acetylcholine (*Chat*- choline acetyltransferase) precursors. Heatmap represents normalized gene counts in TPM. B) *Adrb1^-/-^Adrb2^-/-^*double knockout or wild-type (WT) mice were subject to *S. pneumoniae* pre-exposure and infection. C) Disease burden was greater in *Adrb1^-/-^Adrb2^-/-^*double knockout mice. Unpaired t-test, n=6-7. D) In response to pre-exposure with *S. pneumoniae 19F* and infection with *6303*, *Adrb1^-/-^Adrb2^-/-^*double knockout mice had worse survival. Log-rank (Mantel-Cox) test; n=8. E) *S. pneumoniae* antigen-specific IgG was reduced in the lungs of *Adrb1^-/-^Adrb2^-/-^*double knockout mice. One-way ANOVA with Holm-Sidak post-hoc test; n=4-5. F) B memory B220^+^ CD19^+^ CD38^+^ CD44^+^ MHCII^+^ Tbet^+^ cells were reduced in *Adrb1^-/-^Adrb2^-/-^*double knockout mice. One-way ANOVA with Holm-Sidak post-hoc test; n=4-8. G) B resident memory B220^+^ CD19^+^ CD38^+^ CD44^+^ IgD^-^ CD62l^-^ MHCII^+^ Tbet^+^ cells were reduced in *Adrb1^-/-^Adrb2^-/-^* double knockout mice. One-way ANOVA with Holm-Sidak post-hoc test; n=4-8. H) CD4^+^ Tbet^+^ IFNγ^+^ cells were increased in *Adrb1^-/-^Adrb2^-/-^* double knockout mice. One-way ANOVA with Holm-Sidak post-hoc test; n=4-8. I) IFNγ concentration in the lungs was reduced in *Adrb1^-/-^Adrb2^-/-^*double knockout mice. One-way ANOVA with Holm-Sidak post-hoc test; n=4-6. Two-three independent experiments for each analysis.

To assess the effect of norepinephrine on B cells, we used *Adrb1^-/-^ Adrb2^-/-^* double-knockout mice, which lack β-adrenergic receptors 1 and 2 (Figure 2B). Compared to wildtype mice, *Adrb1^-/-^ Adrb2^-/-^* mice had higher bacterial burden following *S. pneumoniae 19F* pre-exposure and infection (Figure 2C), reduced survival following 19F pre-exposure and *6303* infection (Figure 2D), and reduced *S. pneumoniae* antigen-specific IgG titers (Figure 2E). In concordance with this, the total number of B memory cell types was reduced with receptor deletion both at the early 48 h post-infection and 29 days post-infection time points (Figure 2F, G, S2, S4).

### Loss of β-adrenergic signaling impairs memory B cell formation

Similar to removing the source of norepinephrine with 6-OHDA, adrenergic receptor deletion increased the number of CD4 and CD8 IFNγ-expressing T cells (Figure 2H, S4), yet IFNγ concentration was suppressed in the lung homogenates of double-receptor-knockout mice (Figure 2I). As in the 6-OHDA experiment, neutrophil numbers were higher in *Adrb1^-/-^ Adrb2^-/-^* mice than in wild-type mice (Figure S4). These data support the idea that either norepinephrine inhibition or adrenergic receptor deletion reduces adaptive immunity in response to priming and infection with *S. pneumoniae*. To rule out the role of myeloid cells, we show that myeloid lineage cells are similar between wild-type and *Adrb1^-/-^ Adrb2^-/-^* mice following our model of priming and infection (Figure S5). Furthermore, IL-10 and TNF, which are highly released by macrophages and dendritic cells, were also similar in both sets of mice (Figure S5).

When we probed further to determine the effect of adrenergic receptor deletion on resident B cells, we found that the prominent transcription factor Tbet was suppressed in *Adrb1^-/-^ Adrb2^-/-^* B cells compared with wild-type B cells at early and late time points post-infection (Figure 2F, G). To confirm the effect of norepinephrine on a memory B cell phenotype, we expanded our flow panel and phenotyped various B cell populations in the lung (Figure S6). We confirmed that memory B cell subsets, including activated, marginal-zone-like, and memory B cells, were reduced by adrenergic receptor deletion (Figure 3, S6). We quantified B cell subsets in mediastinal lymph nodes and did not observe significant differences between groups at early and late time points (Figures S6, S7), suggesting that the major effect of norepinephrine occurs within the lung-resident compartment. This supports the notion that norepinephrine binds to B cells and acts to stimulate memory B cells or that norepinephrine induces a memory phenotype(**37, 38**).

**Figure 3.**
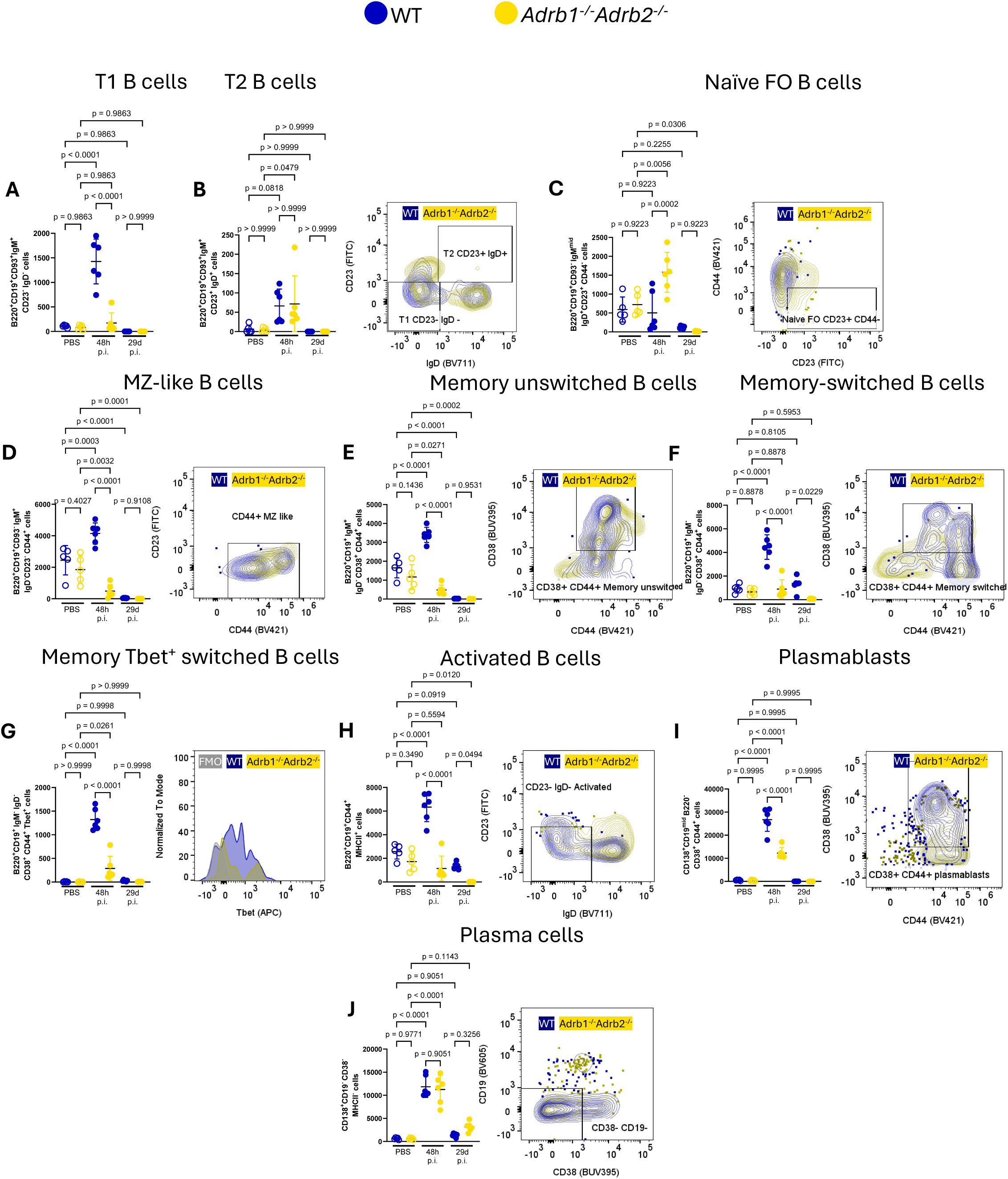
*Adrb1^-/-^Adrb2^-/-^* deletion predominantly affects B cells associated with memory and activation, 48 h and 29 days post-infection. B cells were isolated from WT and *Adrb1^-/-^ Adrb2^-/-^* lungs following our priming and infection model. A) T1 transitional B220^+^ CD19^+^ CD93^+^ IgM^+^ CD23^-^ IgD^-^ cells were greater in wild-type compared to *Adrb1^-/-^Adrb2^-/-^* mice. B) Deletion of β-adrenergic receptors did not affect T2 transitional B220^+^ CD19^+^ CD93^+^ IgM^+^ CD23^+^ IgD^+^ B cells. C) Naïve follicular type B220^+^ CD19^+^ CD93^-^ IgM^mid^, IgD^+^ CD23^+^ CD44^-^ B cells were greater when β-adrenergic receptors were deleted. D) Marginal zone (MZ)-like B220^+^ CD19^+^ CD93^-^ IgM^+^ IgD^-^ CD23^-^ CD44^+^ B cells were greater in wild-type compared to *Adrb1^-/-^Adrb2^-/-^*mice. E) Memory unswitched B220^+^ CD19^+^ IgM^+^ IgD^-^ CD38^+^ CD44^+^ B cells were greater in wild-type compared to *Adrb1^-/-^Adrb2^-/-^* mice. F) Memory-switched B220^+^ CD19^+^ IgM^-^ IgD^-^ CD38^+^ CD44^+^ B cells were greater in wild-type compared to *Adrb1^-/-^Adrb2^-/-^* mice. G) Memory-switched Tbet^+^ B cells were also greater in wild-type compared to *Adrb1^-/-^Adrb2^-/-^* mice. H) Activated B220^+^ CD19^+^ CD44^+^ MHCII^+^ B cells were greater in wild-type compared to *Adrb1^-/-^Adrb2^-/-^* mice. I) Plasmablasts CD138^+^ CD19^mid^ B220^-^ CD38^+^ CD44^+^ cells were greater in wild-type compared to *Adrb1^-/-^Adrb2^-/-^* mice. J) Plasma CD138^+^ CD19^-^ CD38^-^ MHCII^-^ cells were unchanged between wild-type and *Adrb1^-/-^Adrb2^-/-^* mice. One-way ANOVA with Holm-Sidak post-hoc test; n=4-6. Two independent experiments.

### Norepinephrine stimulates T cells and B cells

To confirm that disease outcomes were confined to B cells, we adoptively transferred isolated B cells from wild-type mice and *Adrb1^-/-^ Adrb2^-/-^* mice into *Ighm^-/-^* mice one day before the initial priming with *S. pneumoniae* (Figure 4A). We show that bacterial burden (Figure 4B) and survival (Figure 4C), along with *S. pneumoniae* antigen-specific IgG titers (Figure 4D), in *Ighm-/-* with B cells lacking β-adrenergic receptors phenocopied those seen after 6-OHDA-induced sympathetic neuron depletion and global β-adrenergic receptor knockout.

**Figure 4.**
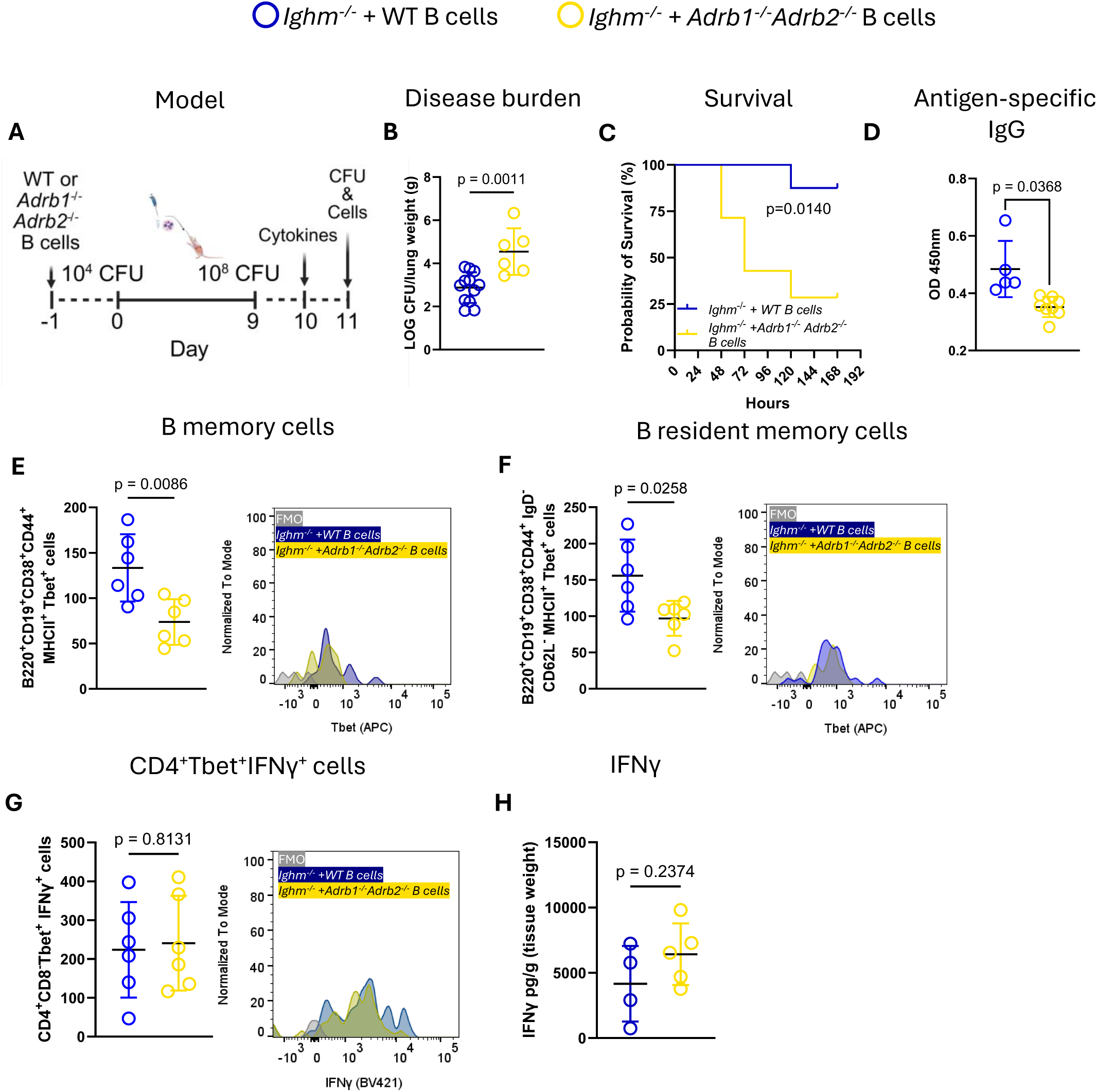
Adoptive transfer of *Adrb1^-/-^Adrb2^-/-^* B cells into *Ighm^-/-^* mice provides B cell-specific deletion of adrenergic receptors, which results in increased disease burden due to a reduction in B cells. A) *Ighm^-/-^* mice received 1x10^7^ *Adrb1^-/-^Adrb2^-/-^*double knockout or wild-type (WT) B cells. Twenty-four hours later, they received the priming dose of *S. pneumoniae*, then on Day 9, received the infectious dose of *S pneumoniae.* B) Disease burden was greater in *Ighm^-/-^*mice with *Adrb1^-/-^Adrb2^-/-^*B cells. Unpaired t-test, n=6-12. C) In response to pre-exposure with *S. pneumoniae 19F* and infection with *6303*, *Ighm^-/-^* mice with *Adrb1^-/-^ Adrb2^-/-^*B cells had worse survival. Log-rank (Mantel-Cox) test; n=7-8. D) *S. pneumoniae* antigen-specific IgG was reduced in the lungs of *Ighm^-/-^* mice with *Adrb1^-/-^Adrb2^-/-^* B cells. Two-way ANOVA with Sidak’s post-hoc test; n=5-8. E) B memory B220^+^ CD19^+^ CD38^+^ CD44^+^ MHCII^+^ Tbet^+^ cells were reduced in *Ighm^-/-^* mice with *Adrb1^-/-^Adrb2^-/-^*B cells. Unpaired t-test; n=6. F) B resident memory B220^+^ CD19^+^ CD38^+^ CD44^+^ IgD^-^ CD62l^-^ MHCII^+^ Tbet^+^ cells were reduced in *Ighm^-/-^* mice with *Adrb1^-/-^Adrb2^-/-^*B cells. Unpaired t-test; n=6. G) CD4^+^ Tbet^+^ IFNγ^+^ cells were increased in *Adrb1^-/-^Adrb2^-/-^*double knockout mice. Unpaired t-test; n=6. H) IFNγ concentration in the lungs was reduced in *Ighm^-/-^*mice with *Adrb1^-/-^Adrb2^-/-^* B cells. Unpaired t-test; n=4-5. Two-three independent experiments for each analysis.

Next, we asked whether the above effects coincided with specific changes in the memory subset of mice that received B cells lacking both β-adrenergic receptors, compared with those that received wild-type B cells. As predicted, infected mice that received *Adrb1^-/-^ Adrb2^-/-^* B cells had fewer Tbet+ memory and resident memory B cell total cell counts than their counterparts that received wild-type B cells (Figure 4E, F, S8). As *Ighm^-/-^* mice have functional T cells, T cell numbers were unaffected by B cell adoptive transfer (Figure 4G, S2, S8), and IFNγ levels were not altered between groups (Figure 4H). Thus, in response to *S. pneumoniae* pre-exposure and infection, norepinephrine affects not only B cells but also T cells, particularly given that both cell types have robust norepinephrine-binding capability (Figure 2A(**33, 34**)).

### Norepinephrine stimulates T cell IFNγ and cooperates with IFNγ to drive B cell IgG

To test how norepinephrine links a sympathetic-T cell-B cell circuit, we used our adoptive transfer system but introduced αCD4- and αCD8-depleting antibodies as previously established(**39**). This produced a comparison between mice with intact T and B cells (*Ighm^-/-^* + isotype + wild-type), depleted T cells and intact B cells (*Ighm^-/-^* + αCD4/CD8 + wild-type), intact T cells and *Adrb1^-/-^ Adrb2^-/-^* B cells (*Ighm^-/-^ +* isotype + *Adrb1^-/-^ Adrb2^-/^* ), and depleted T cells and *Adrb1^-/-^ Adrb2^-/-^* B cells (*Ighm^-/-^ +* αCD4/CD8 + *Adrb1^-/-^ Adrb2^-/-^*; Figure 4A). We found that disease burden was higher in mice in which either T or B cells had been altered (Figure 5B). As expected, IFNγ was reduced with CD4 and CD8 depletion, with similar differences in *S. pneumoniae antigen*-specific IgG titers (Figure 5C, D). To confirm that αCD4- and αCD8-depleting antibodies did, in fact, reduce T cell counts, we analyzed T cells from mice that received isotype or depleting antibodies and demonstrated that T cell counts were reduced with depleting antibodies (Figure 5E).

**Figure 5.**
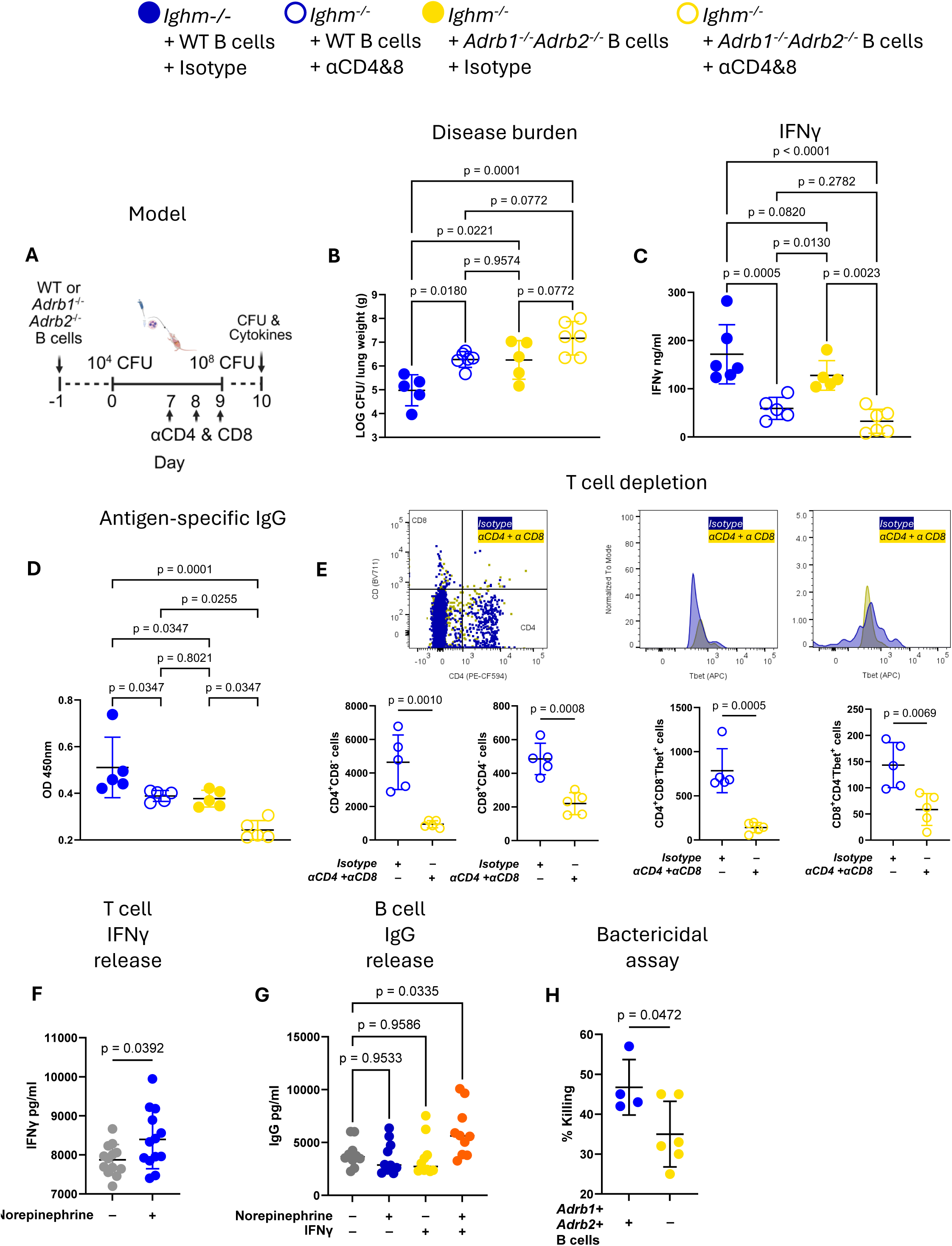
Inhibiting T cells in mice with the adoptive transfer of *Adrb1^-/-^Adrb2^-/-^* B cells in *Ighm^-/-^* mice exacerbates disease burden by a reduction of IFNγ and antigen-specific IgG. A) *Ighm^-/-^*mice received 1x10^7^ *Adrb1^-/-^Adrb2^-/-^*double knockout or wild-type (WT) B cells, then were randomized to receive neutralizing antibodies for CD4 and CD8 or isotype, and then, 24 h later, received the priming dose of *S. pneumoniae 19F* and on Day 9 received the infectious dose of *S. pneumoniae.* B) Disease burden was greater in *Ighm^-/-^*mice with *Adrb1^-/-^Adrb2^-/-^*B cells + isotype and worsened further with the neutralization of CD4 and CD8 T cells in comparison to *Ighm^-/-^,* which received WT B cells + Isotype or WT B cells + CD4 and CD8 neutralizing antibodies. ANOVA with Holm-Sidak post-hoc test, n=5-6. C) IFNγ was reduced with CD4 and CD8 neutralizing antibodies. ANOVA with Holm-Sidak post-hoc test, n=5-6. D) *S. pneumoniae* antigen-specific IgG was reduced in a stepwise manner between groups. Two-way ANOVA with Dunnett’s post-hoc test; n=5-6. E) CD4 and CD8 T cells were analyzed from mice treated with Isotype or neutralizing CD4 and CD8 antibodies. Neutralizing antibodies significantly reduced all T cells and subtypes. n=5 per group, Welch’s unpaired t-test. F) Naïve T cells were isolated from WT spleens and cultured with or without norepinephrine (NE). Norepinephrine stimulated greater T cell IFNγ release. Unpaired t-test; n=13. G) Naïve B cells were isolated from WT spleens and cultured with or without NE, IFNγ, or NE+IFNγ. ANOVA with Dunnett’s post-hoc test. H) *S. pneumoniae 19F* was incubated with decomplemented WT or *Adrb1^-/-^ Adrb2^-/-^*serum. Naïve neutrophils harvested from bone marrow were added, and the remaining bacteria were enumerated following 30 minutes of incubation. Unpaired t-test; n=4-6. Two independent experiments for each analysis.

To confirm this mechanism, we cultured T cells and B cells separately. T cells produced significantly more IFNγ when stimulated with norepinephrine(**32**) (Figure 5F). Meanwhile, when B cells were cultured, both norepinephrine and IFNγ were required to increase IgG production(**40–42**) (Figure 5G). These data support the hypothesis that norepinephrine stimulates T cells to increase IFNγ release, but that the combination of IFNγ and norepinephrine is required for complete B cell stimulation to increase *S. pneumoniae antigen*-specific IgG production(**40–42**).

To confirm that norepinephrine-mediated stimulation of B-cell *S. pneumoniae* antigen-specific IgG release and subsequent bacterial opsonization are necessary for successful bacterial clearance, we conducted an *in vitro* neutrophil bactericidal killing assay. Consistent with our hypothesis, wild-type de-complemented serum enhanced bacterial killing by naïve neutrophils compared to de-complemented serum harvested from *Adrb1^-/-^ Adrb2^-/-^* mice, confirming that norepinephrine affects humoral immunity (Figure 5H). Although neutrophil numbers increased after sympathectomy or β-adrenergic receptor deletion, their bactericidal activity was impaired due to reduced IgG-mediated opsonization. Thus, norepinephrine indirectly governs neutrophil function by enabling antibody-dependent clearance.

## Discussion

Our data support a two-step model in which norepinephrine first stimulates T cells to produce IFNγ, and IFNγ then cooperates with norepinephrine to assist with lung-resident memory B cells. This dual-signal requirement provides a mechanistic explanation for why disruption of either T cell IFNγ or B cell adrenergic signaling impairs IgG production and bacterial clearance.

Multiple neural pathways act to regulate immunity at mucosal sites(**1**). Sensory neurons, along with epithelial cells, are among the first in line to detect pathogens(**19, 20**). We have demonstrated that sensory neurons preferentially release specific neuropeptides, which are dependent on the immunogen but, ultimately, serve to increase immunoglobulin production(**21**). However, nervous system regulation of organ homeostasis can also involve classic autonomic reflex arcs that utilize central nervous system integration and efferent activation of the autonomic nervous system.

The majority of parasympathetic efferent activation involves the release of acetylcholine(**22**). With regard to adaptive immunity, B cells do possess nicotinic receptors(**43**). However, it appears that *chrna7* and *chrna9* stimulation have a suppressive effect on B cells, as germinal center egress is suppressed with vagal stimulation(**43, 44**). Further, no changes to immunoglobulin production were noted with vagal efferent stimulation, which would activate parasympathetic efferent acetylcholine release in the lungs(**44**). Finally, B cells have been shown to produce acetylcholine, and B-cell-derived acetylcholine has a regulatory effect to manage aberrant overproduction of lymphoid cells(**43**). As such, it is possible that vagal-to-vagal reflexes may not directly enhance pulmonary adaptive immunity.

Recent studies have mapped out Th+ neurons in the lung(**23–25**). The main sympathetic neurotransmitter is norepinephrine, of which Th is its precursor. Norepinephrine has been noted to augment immunoglobulin production *in vitro* and in response to injury(**27, 45, 46**). Norepinephrine has also been noted to suppress immunoglobulin production in certain disease-specific contexts(**45, 47, 48**). It is apparent that a temporal and disease-specific context can affect noradrenaline’s effect on immunity(**46, 47**). Our data demonstrate that local sympathetic signaling within the lung regulates T cell and B cell effector functions and stimulates T cell and B cell resident memory cells, thereby enhancing *S. pneumoniae* antigen-specific IgG production, which facilitates bacterial opsonization and the successful clearance of S. *pneumoniae* by neutrophils. We further demonstrate that, with respect to disease outcomes, norepinephrine’s effect on the adaptive response mediates a dual stimulation of T and B cells. Specifically, we provide support that norepinephrine enhances T cell IFNγ release(**32, 45**), and that this effect is likely mediated by canonical pathways, as a lack of norepinephrine or norepinephrine receptors increases intracellular IFNγ in T cells. Subsequently, norepinephrine and IFNγ are both required to increase antigen-specific antibody production(**32, 40–42**).

Norepinephrine either promotes the development of B memory/B resident memory cells by increasing signaling through the Tbet transcription factor, as Tbet is necessary for the establishment of memory(**29, 37, 38**), or specifically stimulates Tbet+ memory B cells. Given that norepinephrine increased T cell IFNγ release, we suggest that co-stimulation of B cells with IFNγ and norepinephrine can shift the memory phenotype. This is consistent with the Immgen database, which shows that memory B cells have increased β2-adrenergic receptor expression(**33, 34**), as well as with previous indications that norepinephrine drives B-cell maturation(**49**), and with the fact that these changes are maintained long-term after infection in our model.

Sympathetic innervation of lymphoid tissues, including the spleen and lymph nodes, appears to terminate predominantly in T cell-rich zones(**49, 50**). Therefore, neural circuits are unlikely to recruit B cells from lymph nodes or secondary lymphoid tissues. Lung tissue-resident B-cells establish themselves early after primary infection(**51, 52**), and are poised to release antibodies(**53, 54**) upon antigen encounter with a challenge infection. It is increasingly recognized that resident B cells establish early(**21, 51, 55**) and do not always require a germinal response to form a mature pool of antibody-secreting cells. Our data demonstrated that CD62L^-^ CD38^+^ CD44^+^ B-resident cells were prominent in our model and were reduced by Th+ neuron ablation or adrenergic receptor deletion. These CD62L^−^ CD38^+^ CD44^+^ Tbet^+^ cells match phenotypes previously described for lung-resident memory B cells that mediate rapid extrafollicular IgG responses(**51, 55–57**). That norepinephrine could augment residency and memory through Tbet fits with recent data showing that Tbet is necessary for B cell maturation(**29, 37, 38**). Moreover, our B cell numbers and phenotypes match closely to the time course of development shown by others(**51, 55**).

Norepinephrine has been shown to suppress neutrophilic recruitment(**58, 59**). Despite an increase in neutrophil numbers with sympathetic ablation or *Adrb1* and *Adrb2* deletion, neutrophils lacking immunoglobulin-mediated pathogenic opsonization cannot effectively clear the pathogen, which is supported by our bactericidal assay Therefore, we suggest that, in an intact system with norepinephrine stimulation of multiple cell types, the sympathetic system serves to restrain undue inflammation by tamping any potential aberrant neutrophilic response, while increasing adaptive immune responses and assisting with the proper function of neutrophilic bacterial clearance.

This study demonstrates that sympathetic efferent neurons play a crucial role in regulating pulmonary humoral immunity and the outcomes of bacterial lung infections. Our data support the idea that β-blockade may increase the risk of community-acquired pneumonia(**60–62**). However, the multiple medications and comorbidities in these patient populations make it difficult to directly attribute any medication to increased risk of infection. Incorporating our previous work(**21**), which shows that sensory neuropeptides stimulate adaptive immunity in the same model of priming and infection, we suggest that norepinephrine acts to direct a specific targeted effect on adaptive immunity by modulating B cell memory. In summary, lung-innervating sympathetic neurons stimulate the release of noradrenaline, which modulates T cells to produce IFNγ, which, alongside norepinephrine, co-stimulates B memory cells to produce *S. pneumoniae* antigen-specific immunoglobulins to assist with bacterial clearance in the lungs.

## Methods

### Mice

All animal experiments were approved by The Lundquist Institute at Harbor UCLA Institutional Animal Care and Use Committee protocol #32183. Mice were housed in a specific-pathogen-free animal facility at The Lundquist Institute. C57BL/6J (Jax: #000664), *Adrb1^-/-^ Adrb2^-/-^* (Adrb1tm1Bkk Adrb2tm1Bkk/J Jax: #003810), *Ighm^-/-^* B6.129S2-Ighmtm1Cgn/J Jax: #002288), mice were purchased from Jackson Laboratories. C57BL/6J wild-type mice were backcrossed for two generations to obtain *Adrb1^+/-^Adrb2^+/−^*mice, which were further bred to generate the littermates for wild-type controls. At the start of experiments, mice were 5-8 weeks old. Age-matched male and female mice were used for experiments.

### Streptococcus pneumoniae culture

*S. pneumoniae* strain 49619 serotype 19F or strain 6303 serotype 3 were purchased from ATCC. Cultures were initially grown on Tryptic Soy Agar (BD Biosciences #236950) supplemented with 5% defibrinated sheep’s blood (Remel #R54012) and cultured for 24h, then re-cultured for an additional 24h in 5% CO_2_ at 37°C. Colonies were then picked for shape and hemolytic activity and cultured in Brain Heart infusion media (BD #DF0037178) in 5% CO_2_ at 37°C. Cultures were adjusted to an OD 600 of 1.0 and resuspended in sterile PBS before inoculation.

### Infection model

*S. pneumoniae* were spun at 4000rpm, 5min, and washed in sterile PBS, then checked to achieve an OD600 of 1.0 and resuspended in sterile PBS. On day 0, mice were infected with 10^4^ CFU of Serotype 19F in 50 µl of sterile PBS via intranasal inoculation. Then, on day 9, mice were infected intranasally with 10^8^ CFU of serotype 19F or 6303 in 50µl of sterile PBS. Cytokines and immunoglobulins were assessed 24 h post-infection. Bacterial burden and cells were assessed 48 h post-infection. Control animals were intranasally infused with 50μL of PBS only(**21**).

### Sympathetic ablation

For lung-targeted sympathetic ablation, 500μg of 6-hydroxydopamine (6-OHDA, dissolved in 50μl of sterile PBS containing 0.1% ascorbic acid) was administered intranasally daily for 3 days, and the animals were rested for an additional 3 days before the first inoculation with *S. pneumoniae* or PBS(**24**).

### Adoptive transfer

B-cells were isolated from spleens of wild-type or *Adrb1^-/-^ Adrb2^-/-^* mice using the EasySep Mouse B-cell negative selection kit (StemCell Technologies #19854). Cells were resuspended in sterile transfer buffer consisting of PBS with 10mM HEPES (Sigma #H3662), 0.5% streptomycin/penicillin (Sigma #P0781), and 2.5%ACDA (SantaCruz Biotechnology #sc-214744). A total of 1x10^7^ cells in 200µl of transfer buffer was injected into the tail vein using a 29-gauge tuberculin syringe into *Ighm^-/-^* mice, 24 h prior to the initial priming with *S. pneumoniae*.

### Bacterial burden determination

Lungs were weighed and then homogenized in 2ml sterile PBS with a tissue homogenizer. Homogenates were serially diluted in sterile PBS and plated on tryptic soy agar plates containing 5% sheep’s blood and 10 µg/ml neomycin (Fisher #BP2669-5). The bacterial CFU were enumerated after overnight incubation at 37°C and 5% CO_2_.

### T cell depletion

For T cell depletion, we followed an established protocol(**39**). Mice were treated with 300μg each of anti-CD4 (GK1.5, BioXCell #BE0003) and anti-CD8 (2.43, BioXCell #BE0061) antibodies intraperitoneally, and with 100μg each of anti-CD4 and anti-CD8 antibodies intranasally on days 7, 8, and 9. Samples were collected on Day 10. Controls received: isotype control (IgG2a, BioXcell #BE0089).

### Tissue collection

For flow cytometry, mice were euthanized by 5% isoflurane inhalation, balance O_2_. The lungs were dissected and perfused transcardially with PBS to remove circulating non-resident cells, coarse dissected, and incubated at 37°C for 45 minutes in 1.75mg/ml collagenase IV (Sigma #C4-22) in PBS; then washed, macerated through a 21-gauge needle, and filtered through a 70µm mesh filter, and then treated for flow cytometry.

For immunoglobulin analysis, lungs were dried, weighed, macerated in 2ml PBS supplemented with 25µl protease inhibitors (consisting of AEBSF HCl (100mM), Aprotinin (80μM), Bestatin (5mM), E-64 (1.5mM), Leupeptin (2mM) and Pepstatin (1mM) Tocris #5500), spun at 4000g for 8min at 4°C and then the supernatant was aliquoted, snap frozen and stored at -80°C until analysis.

### Cytokine, neurotransmitter, and immunoglobulin analysis

Enzyme-linked immunosorbent assay (ELISA) kits were used according to the manufacturer’s instructions, and the plates were analyzed with a BioTek Synergy H1 or BioLegend MiniELISA plate reader. The following commercially available ELISA kits were used: Norepinephrine (Novus Biologicals, #197247), Acetylcholine (Thermo Fisher Scientific, #EEL012) and IFNγ (R&D Systems, #DY485-05), IL-10 (R&D Systems, #DY417-05), TNF (R&D Systems, #DY410-05).

For *S. pneumoniae* antigen*-*specific IgG determination, crude *S. pneumoniae* (12.53µg/ml ATCC #49619) extract was incubated in 96-well plates overnight at 4°C, then blocked with 2%BSA for 2h at RT. Then, log dilutions of lung homogenates (from groups above) were incubated at RT for 2 hours. Goat anti-mouse IgG HRP-conjugated secondary antibody (Abcam #ab205719, 1:1000) was then added and incubated for 2 hours at RT. The ELISA was developed with 0.5% TMB solution (Fisher #AAJ61325AP) and stopped with 2N H_2_SO_4_ (Fisher #828016), and OD450 for each dilution and group was evaluated. Operators were blinded to sample condition.

### Flow cytometry

Red blood cells were lysed with ACK lysing buffer (Gibco #A10492-01), treated with Fc Block (Biolegend TrustainFcX #101320), and resuspended in FACS buffer (HBSS-Gibco #10010-023 with 2% FBS Gibco #10082147). Incubations with antibody cocktails were conducted at 4°C for 60min, and samples were washed twice and resuspended in FACS buffer (HBSS + 2% FBS, Gibco #10082147). For intracellular staining, cells were fixed/permeabilized with the BD Cytofix/CytoPerm Kit (#554714), washed, and stained overnight at 4°C. Flow cytometry was conducted on a Symphony A5 flow cytometer (BD). Data were collected using BD DIVA software, and files were analyzed using FlowJo (Treestar, version 10.0.8r1). A live-cell stain (APC-Cy7, Invitrogen #L34976A) was used to exclude dead cells. Positive staining and gates for each fluorescent marker were defined by comparing complete stain sets with fluorescence minus one (FMO) control stain sets. All antibodies were used at 1:100. Antibodies used: CD45 (FITC Biolegend #103108, clone:30-F11), B220 (APC Biolegend #103212, clone: RA3-6B2), B220 (BV510 Biolegend #103248, clone: RA3-6B2), CD19 (BV605 Biolegend 115540, clone: 6D5), CD23 (FITC Biolegend 101605 clone B3B4), CD93 (PE Biolegend 136503 clone AA4.1), CD38 (BUV395 Invitrogen 363038182, clone:90), CD44 (BV421 Biolegend 1#03040, clone: IM7), CD62L (PE-Cy7 Biolegend 1#04418, clone: MEL-14), IgD (BV711 BD Horizon #564275, clone: 11-26c2a), IgM (BUV805 BD Optibuild #749307, clone: II/41), CD69 (PE Biolegend #104508, clone: H1.2F3), CD138 (BV510 Biolegend #142521, clone: 281-2), CD138 (PE-Cy7 Biolegend 112505 clone W205IE), MHCII (Percp5.5 Biolegend #107626, clone: M5/114.15.2), CD4 (PE/Dazzle594 Biolegend #100456, clone: GK1.5), CD8 (BV711 Biolegend #344734, clone: SK1), RorγT (BV510 BD Horizon #567177, clone: Q31-378) T-bet (APC Biolegend #644814, clone: 4B10), Ly6G (BB700 Biolegend #108428, clone RB6-8C5), CD11b (FITC Biolegend #101205, clone M1/70), CD11c (BV421 Biolegend #117330, clone N418), CD45 (PE-Cy5 Biolegend #103110, clone 30-F11), CD64 (PE Biolgend #139304, clone X54-517.1), F4/80 (BV650 Biolegend #123149, clone BM8). Analysis was run by operators blinded to condition.

Gating strategies.

1. Lymphocytes → Singlets →Live → CD45^+^ →

i. Ly6G^+^ (Neutrophils)
ii. CD138^+^ (Plasma cells)

a. B220^+^ → CD19^+^ → CD38^+^ CD44^+^ (B memory cells)

i. IgD^-^ CD62L^-^ (B resident memory cells)
ii. IgD^-^ IgM^-^ (Isotype switched B cells)
2. Lymphocytes → Singlets → Live → CD45^+^ →

a. B220^+^ → CD19^+^ → CD38^+^ CD44^+^

i. MHCII^+^ → Tbet^+^ (B memory cells)
ii. IgD^-^ CD62L^-^ → MHCII^+^ → Tbet^+^ (B resident memory cells)
iii. IgD^-^ IgM^-^ → MHCII^+^ → Tbet^+^ (Isotype switched B cells)
3. Lymphocytes → Singlets → Live → CD45^+^ →

a. CD4^+^ CD8^-^ →

i. Rorγt^+^ → IFNγ^+^
ii. Tbet^+^ → IFNγ^+^
b. CD8^+^CD4^-^ →

i. Rorγt^+^ → IFNγ^+^
ii. Tbet^+^ → IFNγ^+^
4. Lymphocytes → Singlets → Live →

a. B220^+^ → CD19^+^ →

i. CD44^+^ MHCII^+^ → CD23^-^ IgD^-^ (Activated B cells)
ii. IgM^+^ IgD^-^ → CD38^+^CD44^+^ (Memory unswitched B cells)
iii. IgM^-^ IgD^-^ → CD38^+^ CD44^+^ (Memory switched B cells)
iv. CD93^+^ IgM^+^ →

1. CD23^-^ IgD^-^ (T1 transitional B cells)
2. CD23^+^ IgD^+^ (T2 transitional B cells)
v. CD93^-^ IgM^mid^ →

1. IgD^+^ → CD23^+^ CD44^-^ (Naïve Follicular like B cells)
2. CD93^-^ IgM^++^ →

1. IgD^-^ CD23^-^ → CD44^+^ (Marginal zone like B cells)
vi. CD138 →

i. CD19^mid^ B220^-^ → CD38^+^ CD44^+^ (Plasmablasts)
ii. CD38^-^ CD19^-^ → MHCII^+^ (Plasma cells)
b. Lymphocytes → Singlets → Live →

a. CD45 → CD11b+ CD11c+ → CD64+ F4/80+ → MHCII+ (Myeloid lineage cells)

Spherotech Accucount (#ACFP-70-5) was added to each sample as per the manufacturer’s instructions and used to calculate total cell counts. All flow cytometry data are expressed as total cell counts.

### Immunofluorescence and microscopy

The lungs and aorta were perfused with PBS for immunostaining, then fixed in 4% paraformaldehyde (PFA) in PBS. Lungs were dissected and postfixed overnight in 4% PFA/PBS at 4°C. Lungs were then cleared according to Scott et al(**63**). Briefly, Tissues were washed with TBS 5 × 1 h, then blocked overnight with 4% Normal Goat Serum, 1% Triton X-100, and 5% powdered milk in TBS. Lungs were incubated with streptavidin Alexa 488 (Life Technologies #S-32354), antibodies for Tyrosine hydroxylase (Millipore Sigma #ZMS1033) for 3 days. Lungs were then washed in TBS 5 × 1 h, then overnight at 4°C, followed by treatment with secondary goat-anti-mouse Cy3 (Jackson Immuno #115-165-146). Lungs were then washed in TBS 5 times for 1 h each and post-fixed overnight in PFA to prevent washout of the labeled antibody during optical clearing. Tissues were treated for 1 hour with 50% methanol, then for 2 hours with 100% methanol to dehydrate, and were then immersed in a 1:2 v/v mixture of benzyl alcohol and benzyl benzoate (BABB). Before BABB clearing, the tissue was positioned for later mounting.

3D light-sheet imaging was performed using an UltraMicroscope Blaze (Miltenyi Biotec) operated in Light Speed Mode. Volumetric datasets were acquired using the Multi Color Mosaic Acquisition mode in ImSpector Pro software (version 8.0.1, Miltenyi Biotec). Images were acquired with a dedicated LVBT 4x objective (NA 0.35) at 1.0x zoom, yielding a total magnification of 4.0x and an XY pixel size of 1.62 µm. The camera ROI was 2048 × 2046 pixels, corresponding to a physical field of view of 3328.00 × 3324.75 µm per tile.

Sample illumination was performed with left- and right-sided light sheets using all active sheets and fixed blending. The light-sheet numerical aperture was 0.163, the system NA was 0.35, the light-sheet thickness was 3.9 µm, and the sheet width was set to 100%. Dynamic focus was not used.

Dual-color fluorescence was acquired using a laser combiner with a 5.00-ms camera exposure time:

Channel 1: 488 nm excitation (30% power) with a 525 nm emission filter.
Channel 2: 561 nm excitation (15% power) with a 620 nm emission filter.

Optical sectioning was performed across a 3,682 µm Z range using a 2 µm step size (1842 Z steps). Mosaic imaging was collected using a 2 × 4 tile grid (X × Y) with a 42-pixel overlap, corresponding to 2% overlap in both X and Y. The requested sample ROI was 3540.60 µm × 11352.60 µm. The total acquisition time for this measurement was 648.13s. Following acquisition, the multi-color mosaic datasets were rendered and reviewed using ImarisViewer software (version 10.1.1, Oxford Instruments. Lightsheet images were completed by the Cedars Sinai Biobank and Research Pathology Resource.

Stitched 3D lung image stacks were processed in Python using Napari(**64**). Volumes were read directly with h5py and analyzed at pyramid level 3 (16.0 × 13.0 × 13.0 µm voxels)(**65, 66**).

A lung mask was generated by smoothing and thresholding the FITC channel. Since light-sheet intensity attenuates with depth (2.0–2.6× across each specimen), a smooth background field (Gaussian, σ = 400 µm)(**67**) was estimated within the lung and all subsequent thresholds expressed as ratios to it.

Airway lumen was segmented as the union of two detectors. Small and intermediate airways were found by multiscale Hessian vesselness filtering (Frangi et al., 1998)(**68**) of the inverted image, binarized by hysteresis thresholding with the upper bound from Otsu’s method(**69**). Because vesselness responds as a ring rather than a filled core inside wide lumens, large proximal airways were instead detected as regions below 0.40 × local background and retained by a 40 µm morphological opening, which excludes the dark but thin alveolar parenchyma. The combined mask was skeletonized and converted to a branch graph with skan(**70**); branch caliber was taken from the distance transform and branch length from per-voxel physical step lengths(**66, 67**).

The nerve channel was thresholded by hysteresis relative to its 99.5th percentile and skeletonized. For each nerve voxel, distance to the nearest airway wall was computed as the minimum of (distance to an airway centerline − the local airway radius). Nerve within 30 µm of a wall was assigned to that branch, and mapped nerve length summed per branch. Branches were binned by mean diameter (25 µm bins); nerve length per bin is reported both as a total and normalized per millimeter of airway(**71**).

The pipeline outputs a per-branch table (madlib plot, component identifier, endpoints, length, branch type, intensity statistics, coordinates, diameter, and mapped nerve length/wall-distance with data-availability flags) was produced to assess the degree of Th+ airway innervation. Source code are available at: https://github.com/CypherKn1ght/MouseLung/tree/main.

Aortas were incubated at 4°C in 30% sucrose/PBS for 2 days and stored in 0.1% sodium azide in PBS until cutting. Lungs were embedded in optimal cutting temperature compound, and 14 μm cryosections were cut at -20°C and then blocked for 4 h in PBS containing 10% donkey serum, 2% bovine serum albumin (BSA), and 0.8% Triton X-100. Sections were immune-stained with Th antibodies as above and appropriate secondary antibody as above, washed, and mounted with Prolong Antifade Diamond (Thermofisher #P36961). Aortae were imaged with a Nikon TI Confocal Microscope. Image intensity was analyzed with Nikon EIS Elements software, using fluorescence intensity normalized to the background for the target channel. Operators were blinded to condition.

### Cell culture

Spleens were used to isolate T and B cells under sterile conditions. B-cells were isolated using the EasySep Mouse B-cell separation kit (StemCell Technologies #19854A). B cells were then treated in B cell culture media made using 438mL RPMI 1640 (Gibco #11835-030), 2.5mL 1M HEPES buffer (Sigma #H3662), 5mL GlutaMAX (Thermo Fisher Scientific #35050061), 5mL penicillin/streptomycin stock (Sigma #P0781), 30mL heat-inactivated fetal bovine serum stock (Gibco #A38400-01), and 454μL β-mercaptoethanol (Gibco #21985-023) and vacuum filter sterilized (0.22μm filter)(**72**). Cells were then treated with either media + IL-4 (20ng/ml, Biolegend #574304) + LPS (10µg/ml, #LS25-1MG Sigma Aldrich), media+IL-4+LPS+IFNγ (10ng/mL, Thermofisher, #31505100UG) IL-4+LPS+ norepinephrine (10ng/ml, Millipore Sigma, #1468501-125MG) or IL-4+LPS+IFNγ+norepinephrine . After 96 hours, the media were stored at -80°C until IgG analysis.

T cells were isolated with the Mojosort CD3 T cell isolation kit (Biolegend, #480024). For T cell culture, 48-well plates were pre-coated with an α-CD3e antibody (1 µg/mL, BioLegend, #100301) for 2 hours at 37°C. Following isolation, T cells were seeded at 5x10^5^ in 0.5ml media consisting of RPMI 1640 (445ml), penicillin/streptomycin (5ml), 1M β-mercaptoethanol (25µl), fetal bovine serum stock (10%, Gibco #A38400-01), αCD28 antibody (1µg/ml, Biolegend, #117003), IL-2 (5ng/ml, Millipore Sigma #I0523-20UG), IL-12 (10ng/ml, Biolegend #573102) with or without norepinephrine (10ng/ml). After 96 hours, the media were stored at -80°C until IFNγ analysis.

### Neutrophil bactericidal assay

Mouse neutrophils were purified as described previously(**21, 73**). In brief, bone marrow cells were flushed from the femurs and tibias of 8-week-old C57BL/6J mice using sterile RPMI 1640 medium supplemented with 10% FBS and 2 mM EDTA and collected into a 50mL screw-top Falcon tube fitted with a 70μm filter. Mouse neutrophils were purified from bone marrow cells (BioLegend MoJo Sort #480057) according to the manufacturer’s instructions. Bone marrow-enriched neutrophils had >98% purity and >93% viability. Neutrophil killing of S*. pneumoniae* 19F was determined by CFU enumeration. *S. pneumoniae* 19F was incubated with 50% decomplemented (20min; 65°C) serum in PBS from wild-type and *Adrb1-/-Adrb2-/-* mice for 30min on ice. Bacteria were washed with PBS, and 1x10^3^ serum-coated *S. pneumoniae* 19F were incubated with 1x10^5^ BM-neutrophils for 30min. Neutrophils were lysed with ice-cold water for 5 minutes, diluted, and the remaining bacterial cells were quantified by culture.

### Sample size and statistical analysis

All replicates, or animal numbers, are detailed in the figure legends. No specific sex effects were visualized; therefore, group differences were analyzed with an unpaired t-test, one-way ANOVA with either Holm-Sidak or Dunnett’s post hoc test. Data were plotted in Prism (GraphPad Version #10.6.1).

## Acknowledgments

This project was supported by NIH grant R01HL17646301-A1 and UCLA CTSI UL1TR001881-01. The content is solely the responsibility of the authors and does not necessarily represent the official views of the National Institutes of Health. Figures 1A, 2B, 4A and 5A and the graphical abstract were made using Biorender under a CC-BY license. The authors gratefully acknowledge the Light Microscopy & Image Analytics Team of the Biobank and Research Pathology Resource in Cedars-Sinai Health Sciences University for their expert assistance with lightsheet microscopy image acquisition and image analysis.

## Author contributions

F.Z. K.D. and, S.A., contributed to the experimental design, conducted the experiments, and analyzed the data. J.S. conducted experiments and analyzed data. R.P, D.A. and, N.K. contributed to data analysis. R.P. analyzed 3D images and generated source code. O.A., S.S., and P.J. assisted with experimental design and analysis. M.S contributed to study design and manuscript preparation. N.J. designed the study, conducted experiments, analyzed data, prepared the manuscript, and figures. All authors agree with the manuscript.

M.S., D.A., and N.J. declare the following competing interests. U.S. Patent Application Serial Number PCT/IB2024/060224. Methods of using sensory neuron neurotransmitters to enhance humoral immunity. The remaining authors do not declare competing interests.

## Definitions

*Adrb1-* β_1_-adrenergic receptor

*Adrb2-* β_2_-adrenergic receptor

IFNγ- interferon gamma

Ighm^-/-^ - µMT mice lack mature B cells

6-OHDA- 6hydroxydopamine (depletes norepinephrine, rendering sympathetic neurons inert)

Tbet- T box expressed in T cells

Th+ tyrosine hydroxylase + (key precursor to the sympathetic neurotransmitter, norepinephrine)

