## Supplementary figures for "Local sympathetic neurons regulate adaptive immunity in response to Streptococcus pneumoniae infection by modulating T- and B-cell effector functions"

Figure S1.

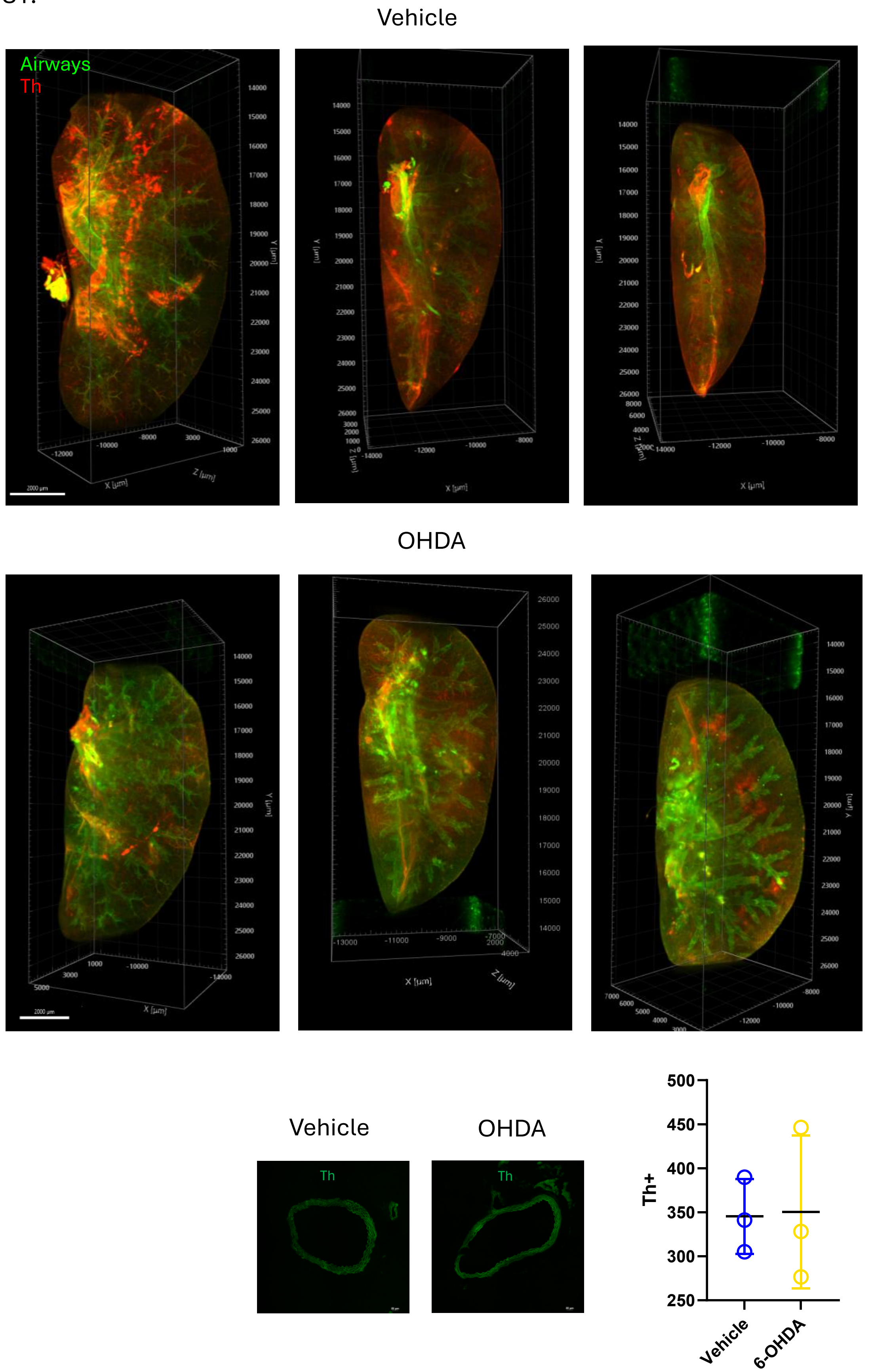


**Figure S1. Tyrosine hydroxylase-positive cells were preserved outside the lungs.** Lightsheet images of cleared lungs from vehicle and 6-OHDA mice.

Intranasal 6-OHDA did not affect aortic innervation by tyrosine hydroxylase + (Th) neurons. Unpaired t-test; n=3.

Figure S2.

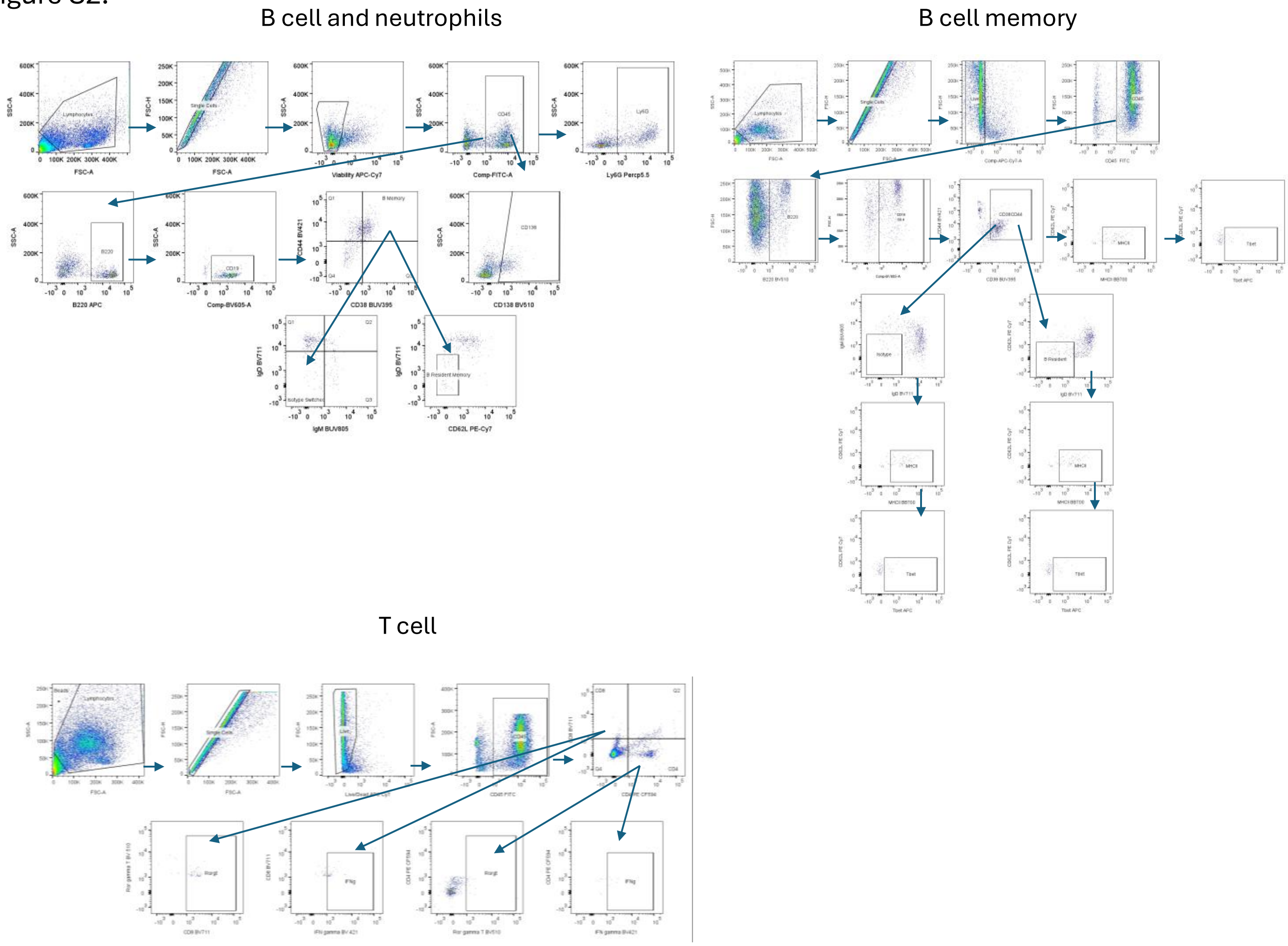


**Figure S2. Gating strategy used for B cells, neutrophils, memory B cells, and T cells.** Gating used for in vivo flow cytometry data from lungs collected on BD Symphony A5.

Figure S3.

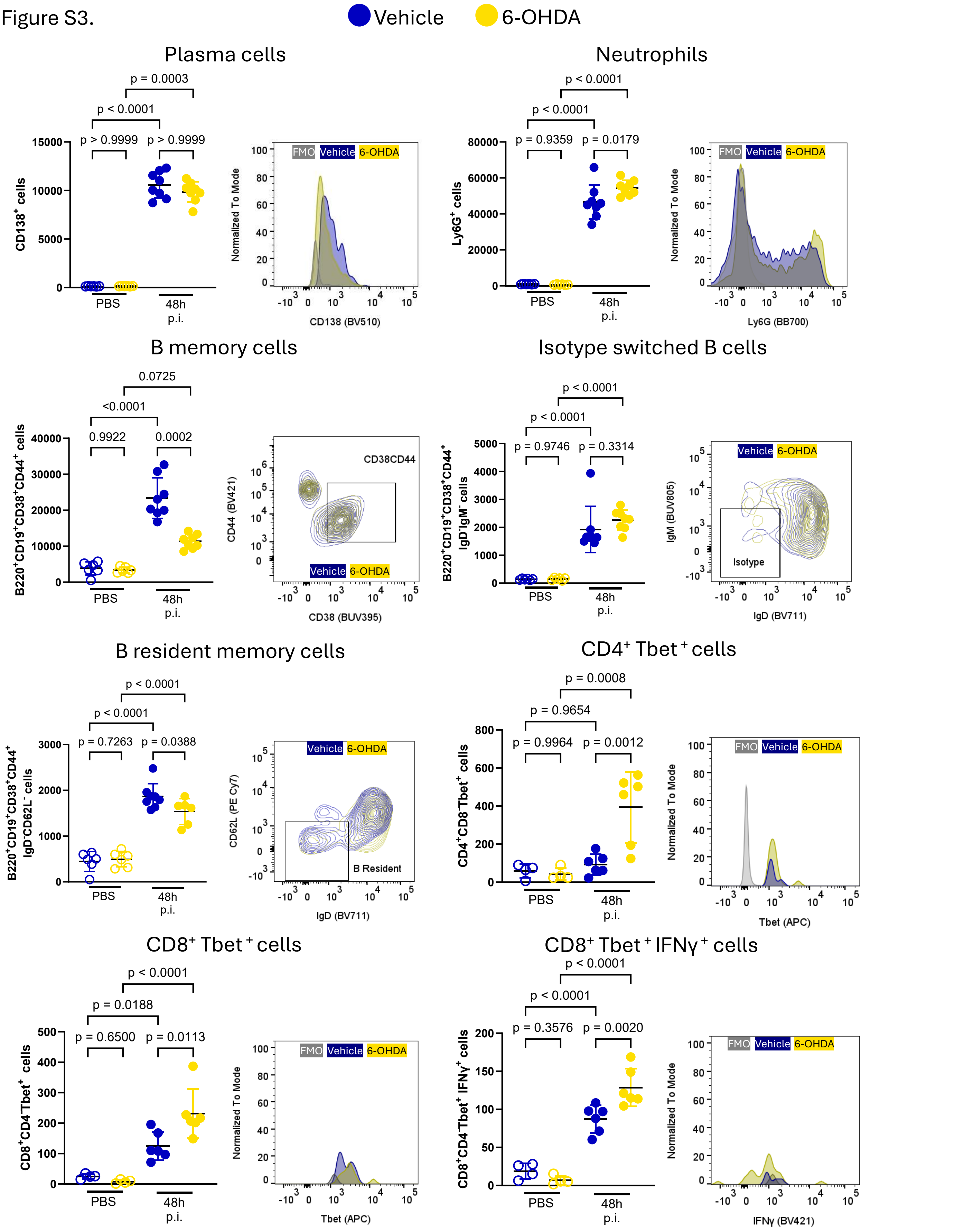


**Figure S3. Lung cells were analyzed from vehicle- and 6-OHDA-treated mice following *S pneumoniae* pre-exposure and infection.** Plasma cells, neutrophils, B memory, isotype switched, B resident memory, Th1, Cytotoxic T cells were analyzed from naïve and infected mice. One-way ANOVA with Holm-Sidak post-hoc test; n=4-8.

Figure S4.

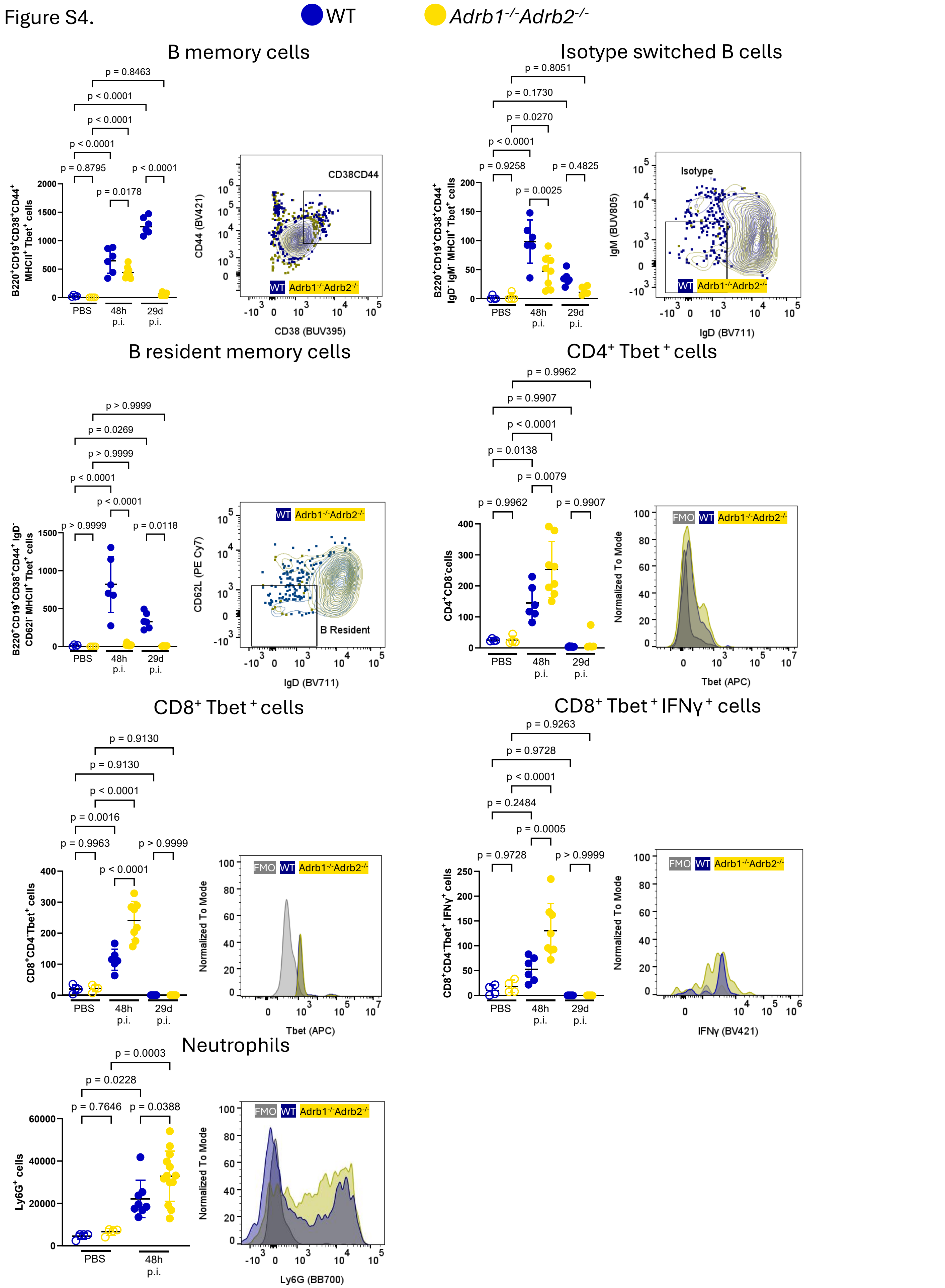


**Figure S4. Lung cells were analyzed from wild-type and *Adrb1$^{-/-}$Adrb2$^{-/-}$* double-knockout mice following S. *pneumoniae* pre-exposure and infection.** B memory, isotype switched, B resident memory, Th1, Cytotoxic T cells, and neutrophils were analyzed from naïve and infected mice. One-way ANOVA with Holm-Sidak post-hoc test; n=4-8.

Figure S5.

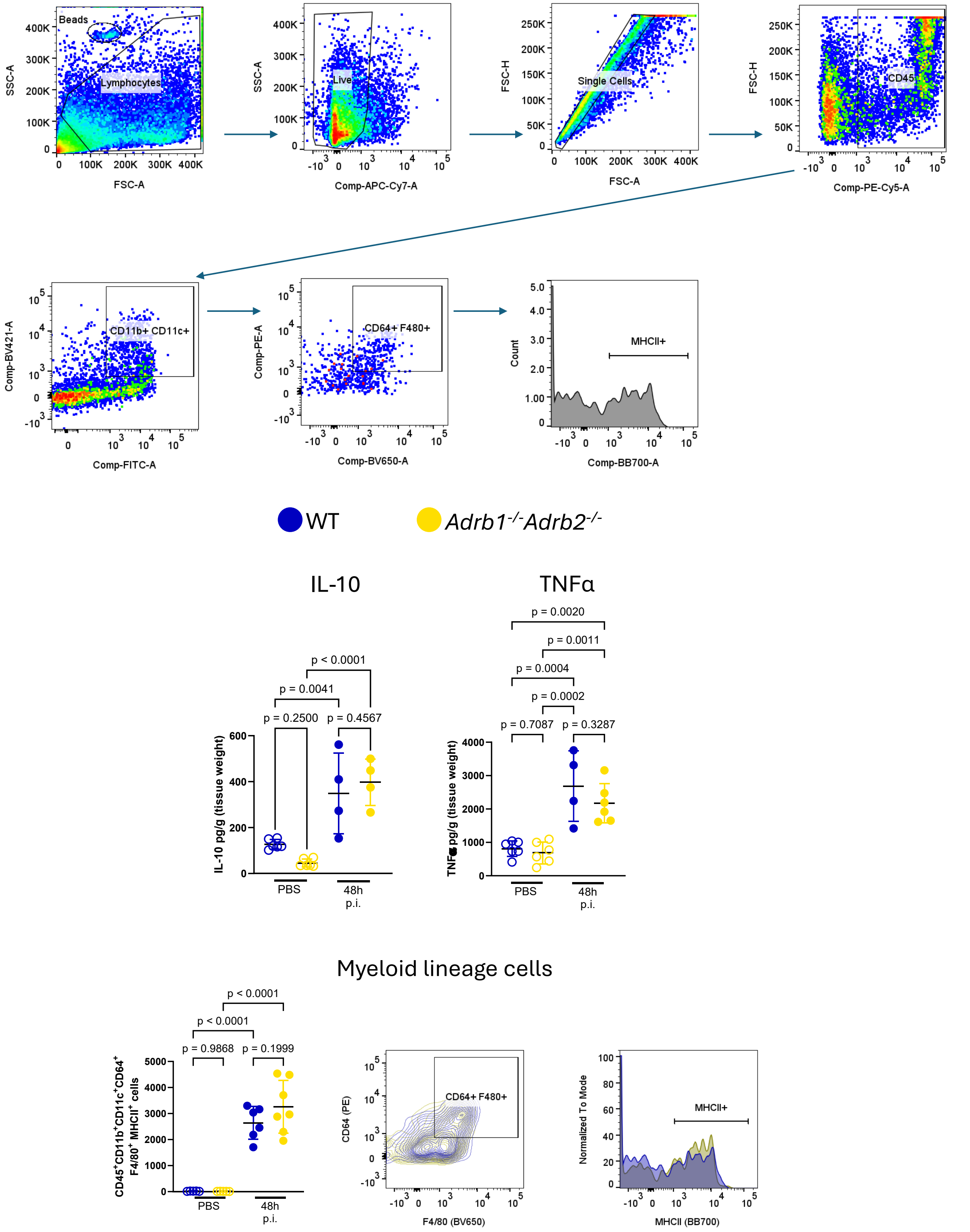


**Figure S5. Myeloid lineage cells and cytokines are not different in wild type and *Adrb1*$^{-/-}$ *Adrb2*$^{-/-}$ mice post-infection.** Gating used for in vivo flow cytometry data from lungs collected on BD Symphony A5 to identify myeloid lineage cells. IL-10 and TNFα were measured from lung homogenates. Myeloid lineage cells (CD45$^{+}$ CD11b$^{+}$ CD11c$^{+}$ CD64$^{+}$ F4/80$^{+}$ MHCII$^{+}$; gated on live singlets) were similar between WT and *Adrb1*$^{-/-}$*Adrb2*$^{-/-}$ naïve mice and after infection. One-way ANOVA with Holm-Sidak post-hoc test; n=4-8.

Figure S6.

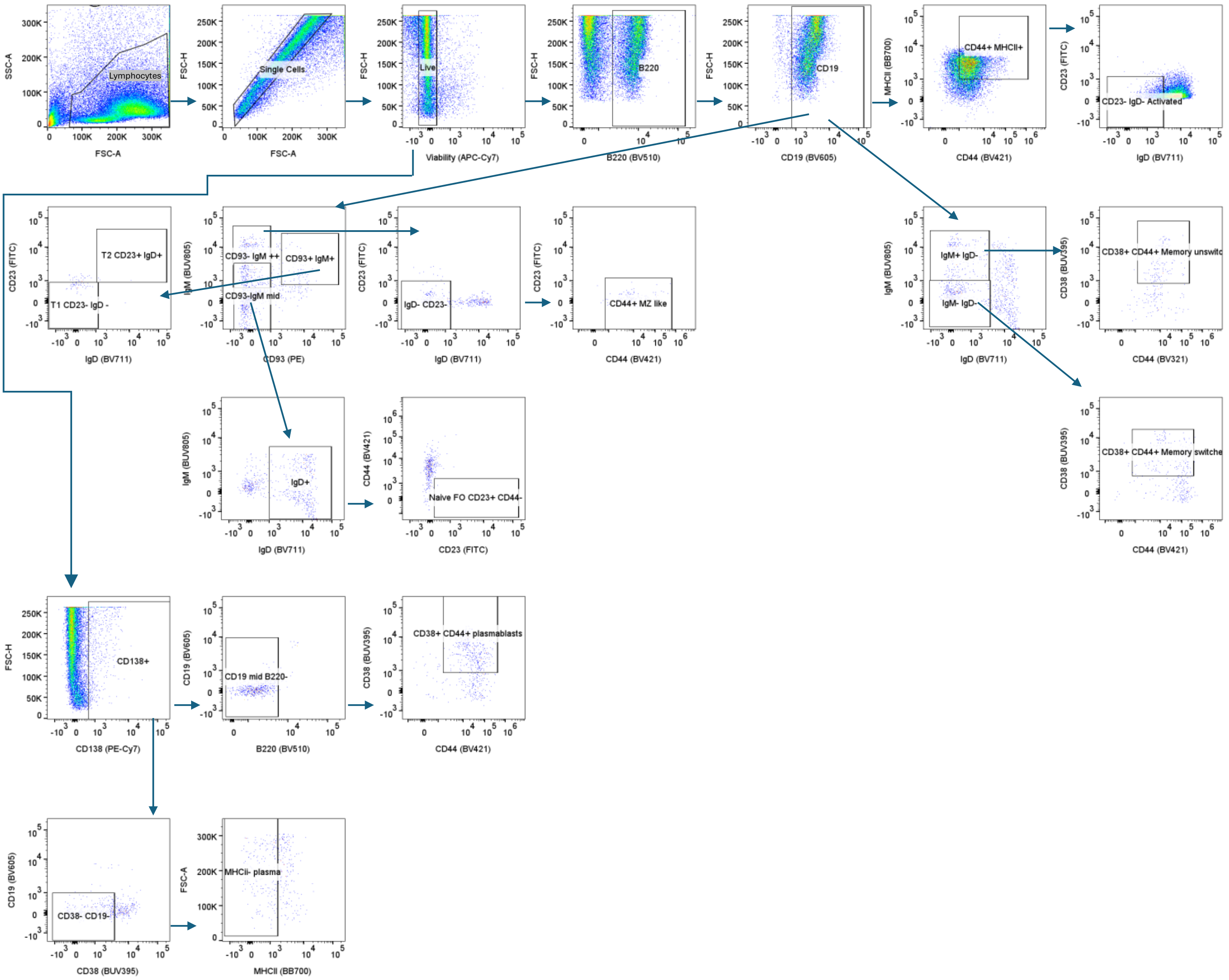


**Figure S6. Gating strategy used for extended B cell phenotyping.** Gating used for in vivo flow cytometry data from lungs collected on BD Symphony A5.

Figure S7.

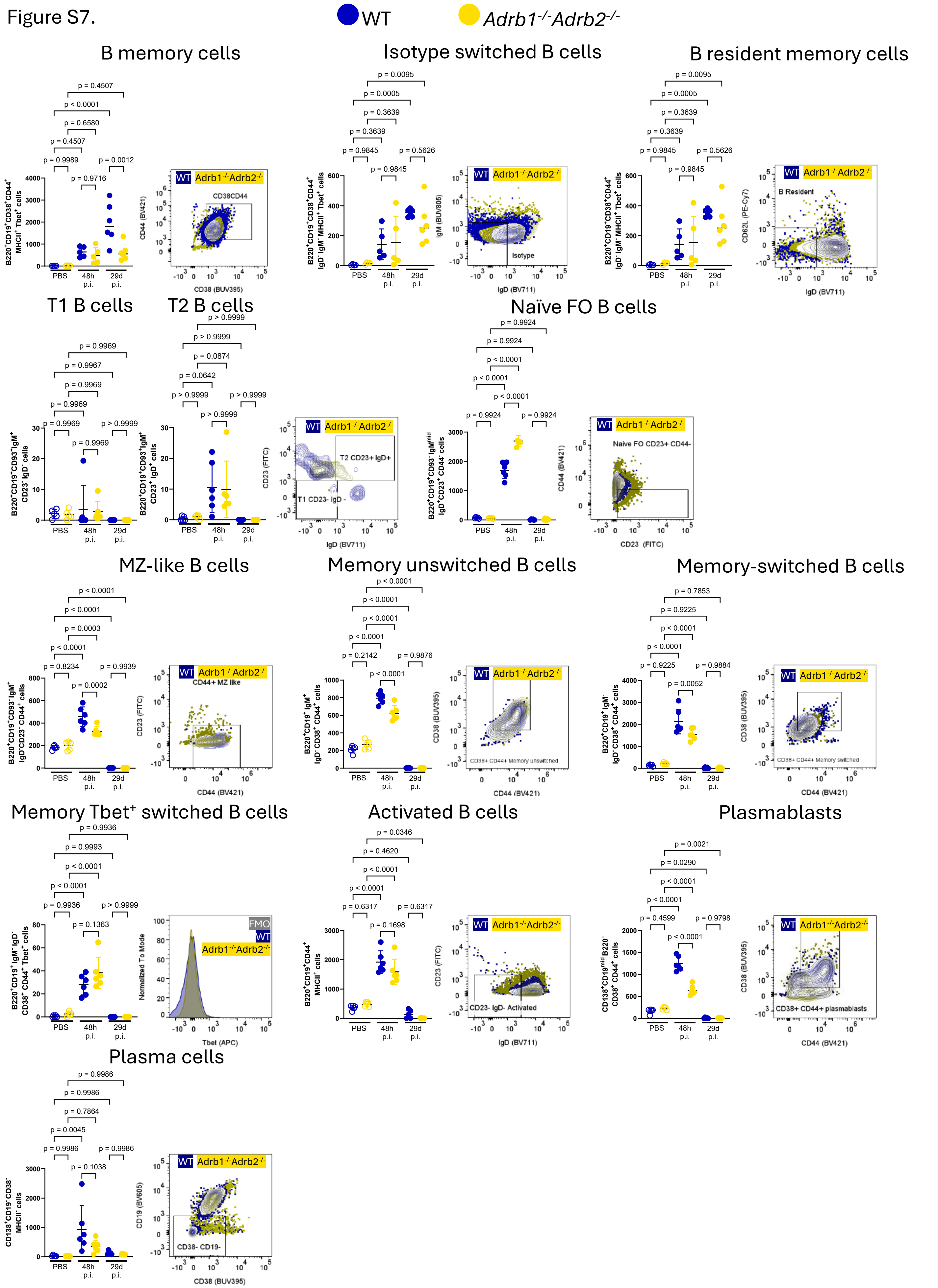


**Figure S7. Cells from lung-draining mediastinal lymph nodes following S. *pneumoniae* pre-exposure and infection.** Cells from the mediastinal draining lymph node were analyzed from naïve and infected mice 48h and 29 days post infection. One-way ANOVA with Holm-Sidak post-hoc test; n=4-8.

Figure S8.

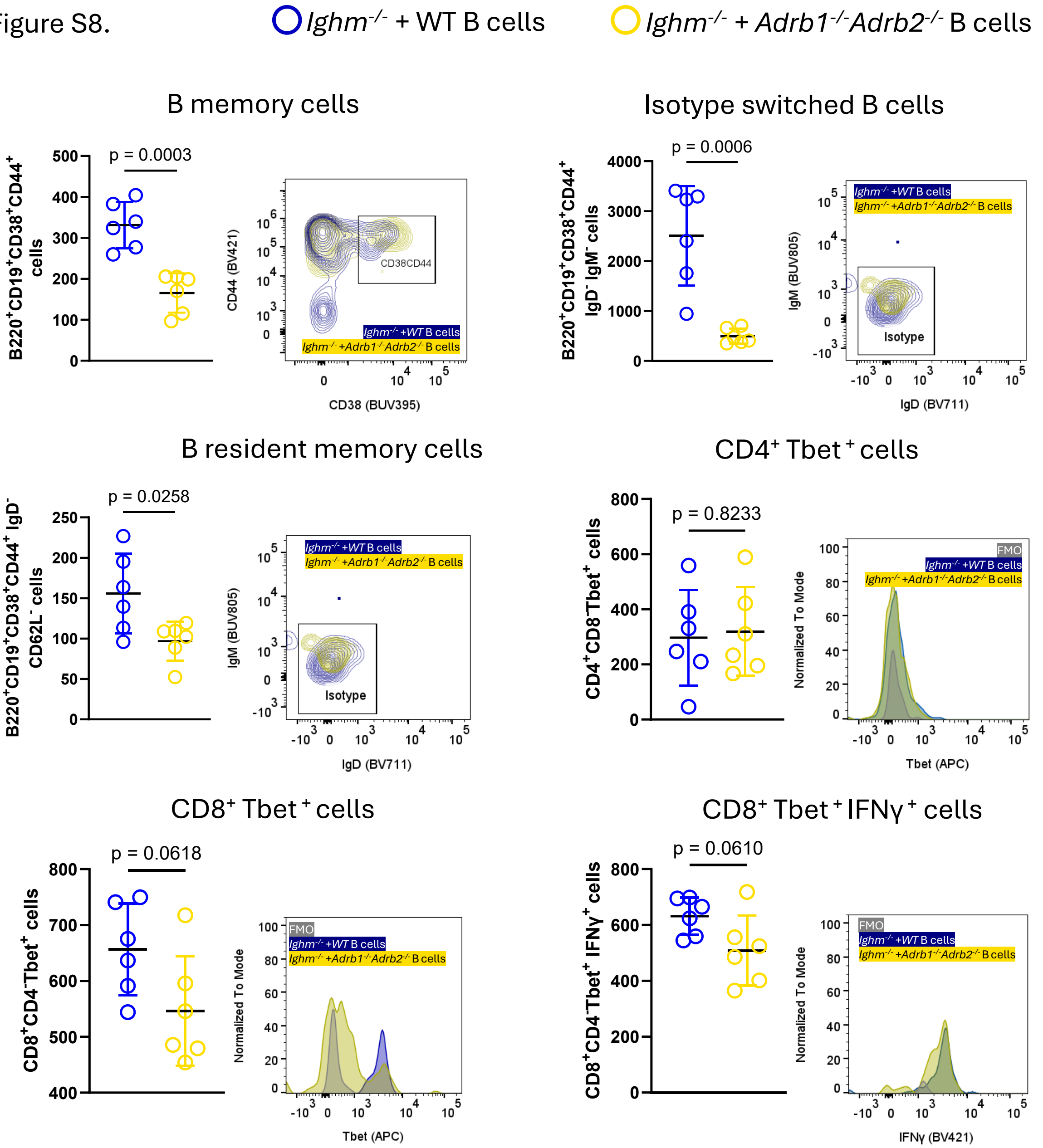


**Figure S8. Lung cells were analyzed in Ighm$^{-/-}$ with either wild-type or *Adrb1$^{-/-}$Adrb2$^{-/-}$* double-knockout B cells following *S pneumoniae* pre-exposure and infection.** Plasma cells, neutrophils, B memory, isotype switched, B resident memory, Th1, Cytotoxic T cells were analyzed from naïve and infected mice. Unpaired t-test.
